# Interpretable Deep Learning Reveals Biologically Relevant Spatial Gene Expression Patterns in Lung Tumors and their Microenvironment

**DOI:** 10.1101/2025.09.17.674367

**Authors:** Vibha R. Rao, Adrienne A. Workman, Liang Lu, Xiaoying Liu, Shrey S. Sukhadia

## Abstract

Lung adenocarcinoma (LUAD), the most common subtype of non–small cell lung cancer (NSCLC) exhibits profound histological and molecular heterogeneity, hindering accurate prognosis and effective treatment. Current approaches to assess this heterogeneity, such as histopathology, molecular profiling, and spatial transcriptomics are constrained by high costs, long turnaround times, and limited tissue availability, making them challenging for widespread prognostic use. To address this gap, we developed XpressO-Lung, an explanatory deep learning model that predicts gene expression heterogeneity, spatially, in tumor and its microenvironment, on hematoxylin and eosin (H&E) based diagnostic (Dx) whole-slide images (WSIs) by learning associations between tissue morphology and the corresponding bulk-transcriptomic data. Utilizing 200 LUAD cases from The Cancer Genome Atlas (TCGA), XpressO-Lung predicted spatial expression patterns of *NAPSA, SLC47A1, TP53I3, KLRB1*, *FAM189A1, TICAM1, CD8A, CXCL13, TTF, CDH3*, *KRT7* and *CDKN2A* genes (biomarkers) on the respective Dx-WSIs with AUCs ranging up to 0.92. More importantly, the predicted spatial gene expression patterns aligned with the known morphologic interactions of the tumor and its microenvironment, capturing biological events directly on Dx-WSIs. By coupling predictive performance with spatial interpretability of gene expression on Dx-WSIs, XpressO-Lung bridges histopathology and bulk-transcriptomics, enabling explainable morpho-genomic analyses to advance biomarker discovery and offer prognostic insights to inform precision oncology in LUAD, especially in low-resource settings.

## Introduction

Lung adenocarcinoma (LUAD) is the most prevalent histological subtype of non-small cell lung cancer (NSCLC), accounting for approximately 40% of all lung cancer diagnoses ^1,2^. Originating from glandular epithelial cells in the distal airways, LUAD is characterized by considerable histological and molecular heterogeneity, often presenting with distinct architectural growth patterns such as lepidic, acinar, papillary, micropapillary, complex glandular, and solid subtypes.^3^. This heterogeneity extends beyond the malignant epithelial cells to include the surrounding tumor microenvironment (TME), which comprises stromal, immune, and vascular elements that significantly influence tumor progression, treatment response and prognosis ^3^. Genomic diversity is also prominent, with frequent oncogenic driver mutations in genes such as *EGFR, KRAS, ALK, STK11,* and *TP53* ^4–6^. Current guidelines, including the WHO and CAP lung cancer biomarker reporting protocols, recommend reflex testing for ALK and ROS1 rearrangements, while the NCCN and IASLC-CAP-AMP guidelines further advocate broad molecular profiling to include EGFR, KRAS, BRAF, MET, RET, NTRK, and PD-L1 as part of routine diagnostic workup^7^.These alterations inform therapeutic strategies, particularly the use of tyrosine kinase inhibitors (TKIs) and immune checkpoint inhibitors, both of which have substantially improved outcomes for subsets of LUAD patients ^8^. Nevertheless, a substantial subset of LUAD cases show unpredictable responses to therapy and widely varying prognoses, underscoring the need for more comprehensive molecular characterization ^9^.

A major obstacle to improving prognostic accuracy in LUAD is the high degree of inter- and intra-tumoral heterogeneity, which includes variation in both cancer cell states and TME composition. Even among patients with identical histological subtypes or driver mutations, differences in gene expression programs can significantly influence tumor progression, immune interactions, and treatment response ^10^. Addressing this complexity requires an expanded genetic landscape and scalable tools capable of capturing transcriptomic diversity across LUAD cohorts.

Spatial transcriptomics technologies have emerged as powerful tools for mapping gene expression across tumor sections, enabling in situ profiling of both cancer and non-cancer compartments with high resolution ^11^. However, their technical demands, high costs, and limited clinical availability pose significant barriers to widespread implementation. Moreover, these methods often require specialized protocols and non-standard tissue preservation techniques, limiting their applicability to large-scale or retrospective prognostic studies ^12^. As a result, there is growing interest in computational approaches that can infer molecular features from more accessible sources, such as routine histopathology slides.

Hematoxylin and eosin (H&E)-stained diagnostic (Dx) whole slide images (WSIs) offer a rich but underutilized source of phenotypic data. Recent developments in computational pathology have leveraged deep learning (DL) models to extract biologically relevant features from WSIs, demonstrating the ability to classify LUAD subtypes ^13^, predict driver mutations ^14^, infer tumor microenvironment composition ^15^, and even approximate transcriptomic signatures ^16^. These models, often based on convolutional neural networks (CNNs) or attention-based architectures, have shown strong performance across aforementioned predictive tasks. However, a persistent challenge in these approaches is their lack of interpretability ^17^.

Most DL models in computational pathology function as “black boxes,” providing slide-level predictions without clear explanations of how these decisions are made or which tissue regions contribute most to the outcome ^18,19^. This lack of interpretability remains a major barrier to its clinical application, particularly for prognostic tasks, as it hinders trust in models’ predictions and complicates biological validation^18^. In the context of prediction of spatial gene expression that drives tumor progression and patient outcomes, the inability to interpret histomorphologic regions along with the predicted spatial gene expression in both tumor and TME, downgrades the utility of these models for prognostic assessment and clinical decision-making for cancer patients. The importance of interpretability in clinical applications has also been recognized at the policy level, with the White House’s National AI Action Plan designating interpretability as a national research priority and emphasizing its critical role in ensuring safe, trustworthy, and clinically meaningful AI deployment in healthcare ^20^.

To address these challenges, we introduce XpressO-Lung, an explanatory deep learning model that predicts spatial gene expression heterogeneity in tumors and TME from H&E-stained Dx WSIs by learning associations between tissue morphology and bulk transcriptomic profiles using our XpressO pipeline ^21^. XpressO-Lung builds upon our prior work in breast ^21^ and melanoma ^22^ cancers, extending the same modular, transparent framework to LUAD. The XpressO pipeline employs a weakly supervised attention-based multiple instance learning (MIL) approach, generating gene-specific spatial heatmaps that highlight high- and low-attention regions of gene expression across both tumor and its microenvironment ^21^. We then analyse these heatmaps by comparing them with the pathologists’ annotations and interpret specific tissue regions driving the prediction of each gene (biomarker), thereby uncovering biologically meaningful spatial biomarker expression patterns within the tumor and TME, that highlight prognostic differences in LUAD. Therefore, XpressO-Lung not only offers the predictive power of state-of-the-art DL models, but also defines a clear methodology for leveraging and interpreting such models to reveal spatial morpho-genomic relationships in cancer, enabling the level of interpretability necessary for trustworthy prognostic application in the clinic.

## Materials and Methods

### Data Collection and Preprocessing

We used an explainable deep learning pipeline ‘XpressO’ ^21^ to build the XpressO-Lung model to predict tissue-based gene expression heterogeneity from the corresponding Dx-WSIs in a weakly supervised and interpretable manner. XpressO spatially connects tissue morphology with gene expression patterns, enabling us to discern morphologic patterns, such as co- or alternate expression of genes in both tumor and TME. It incorporates four core modules: Segmentation, Feature Extraction, Classification, and Heatmap that enable seamless deployment across diverse cancer types with minimal reconfiguration, while effectively capturing spatially informative features from both tumor and TME regions. Dx-WSIs of LUAD from 200 patients were obtained from The Cancer Genome Atlas (TCGA-LUAD) via the Genomic Data Commons (GDC) portal^23^. These WSIs were processed using the segmentation module in XpressO ^21^ which leverages the CLAM framework ^24^ to identify tumor-enriched regions by assigning attention scores to image patches and selecting the most informative regions for analysis. The WSIs were tiled into non-overlapping 256×256 pixel patches at 20× magnification ^25^. The resulting patches were passed through the feature extraction module in XpressO ^21^, which uses a pretrained Vision Transformer (ViT-L/16) ^26^ called Unified Network for Instance-level Representation Learning (UNI) ^27^, trained with DINOv2 ^28^ to extract high-dimensional feature embeddings optimized for histopathological image analysis. These features were then retained for downstream classification.

### Collection and Processing of RNA-seq Data

Corresponding bulk RNA sequencing (RNA-seq) data for the same 200 LUAD patients were downloaded from the TCGA-LUAD portal in the form of fragments per kilobase of transcript per million mapped reads (FPKM). To prepare the dataset for gene expression classification, FPKM values were binarized into “high” and “low” categories using XpressO’s custom binarization script ^21^. For each gene, the median expression value across all patients was used as the binarization threshold. Samples with expression values greater than or equal to the median were labelled as high-expression (“1”), while those below the median were labelled as low-expression (“0”).

Following binarization, the dataset was split into training, validation, and testing subsets using a predefined XpressO script. A range of k values (k = 4 to 14) was used to create k-folds for each gene. The value of k varied by gene (e.g., k = 10 or 13), resulting in different training, validation, and testing ratios (e.g., 80:10:10 [training:validation:testing] for k = 10; 74:13:13 for k = 13).

Binary expression labels were assigned at the slide level, allowing each WSI to be categorized as high (“1”) or low (“0”) expression for a given gene. These labelled subsets were then used to train and evaluate the deep learning model for gene expression prediction using WSIs

### Model Training

Gene expression prediction was performed using the classification module in XpressO, which is based on the CLAM-SB architecture ^24^. For each gene, a weakly supervised model was trained to classify binarized expression status using the top-ranked patch embeddings from each tissue-slide in the training set. Training was conducted using binary cross-entropy loss, with the Adam optimizer (learning rate = 2×10^−4^) over 200 epochs. The model was trained to weigh instance-level features using attention-based pooling and output a slide-level prediction for each gene’s expression class.

### Model Evaluation and Metrics

The trained model was evaluated on independent test sets for each gene of interest based on the data-split corresponding to the best-performing k-fold during training. For each gene, the model that achieved the best validation performance during cross-validation was selected, and its performance was then evaluated on the corresponding held-out (or independent) testing set (i.e. 10% of the total set, in case of k=10-as an example). Evaluation metrics included area under the receiver operating characteristic curve (AUC-ROC), accuracy, precision, recall and F1-score. These metrics were computed separately for each gene based on the performance of the model on the independent test set.

### Visualization of the Distribution of High and Low Gene Expression on WSIs

Interpretation of spatial heatmaps of gene expression on WSIs were conducted using XpressO’s HeatMap module wherein the attention scores for each patch of the test WSI got mapped to their corresponding spatial coordinates. These heatmaps aided visualization of spatial patterns associated with high versus low expression and co-versus alternating expression of gene pairs on WSI, allowing for the identification of histologic features contributing to the patterns of expression across multiple genes using gene-specific spatial heatmaps from the same test WSI clubbed together for a paired interpretation. Predicted heatmaps were compared with pathologists’ annotations to evaluate alignment between model-derived predictions and established histomorphologic patterns.

## Results

### Genes of interest

Analysis of bulk RNA-seq profiles from 200 LUAD tumors in the TCGA cohort revealed a broad range of gene expression patterns across patients. Twelve genes were selected based on prior literature implicating their varying expression in tumor and TME in LUAD: *CD8A*, *CDH3*, *CDKN2A*, *CXCL13*, *FAM189A1*, *KRT7*, *NAPSA*, *SLC47A1*, *TICAM1*, *TP53I3*, *TTF1*, and *KLRB1* (Table 1a). Their fpkm values are depicted in Table 1b. These genes encompass diverse facets of LUAD pathogenesis, including tumor lineage specification, cell cycle regulation, immune infiltration, epithelial plasticity, and therapeutic response. *TTF1* (NKX2-1) ^29–32^, *NAPSA* (Napsin A) ^33–37^, and *KRT7* ^38–42^ are canonical immunohistochemical (IHC) markers routinely used to distinguish LUAD from squamous and metastatic tumors. *CD8A* ^43–48^ and *KLRB1* (CD161) ^49–51^ reflect cytotoxic lymphocyte infiltration and immune competence, while *CXCL13* ^52–57^ organizes tertiary lymphoid structures and predicts favorable response to immunotherapy.

**Table 1a:**
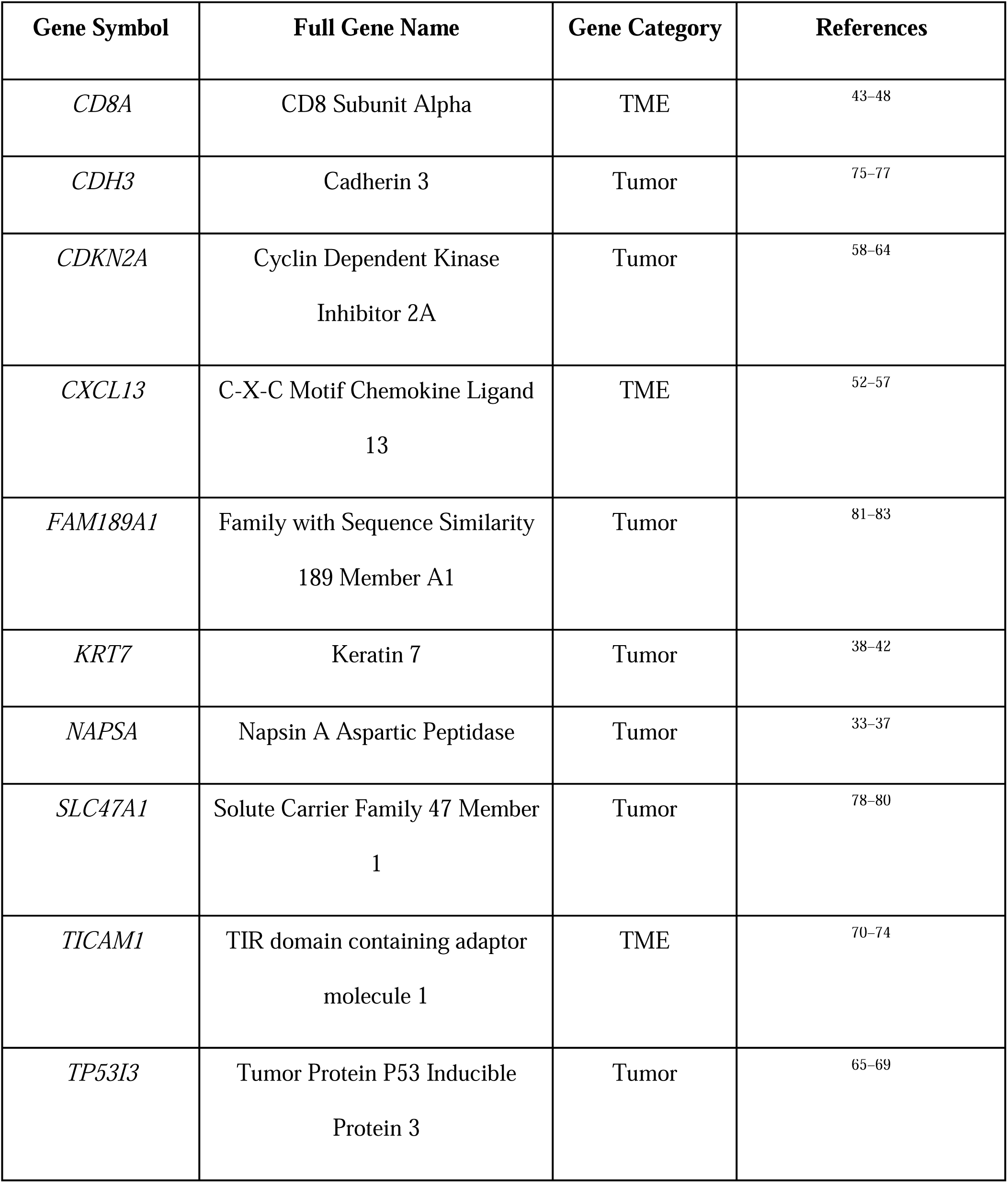
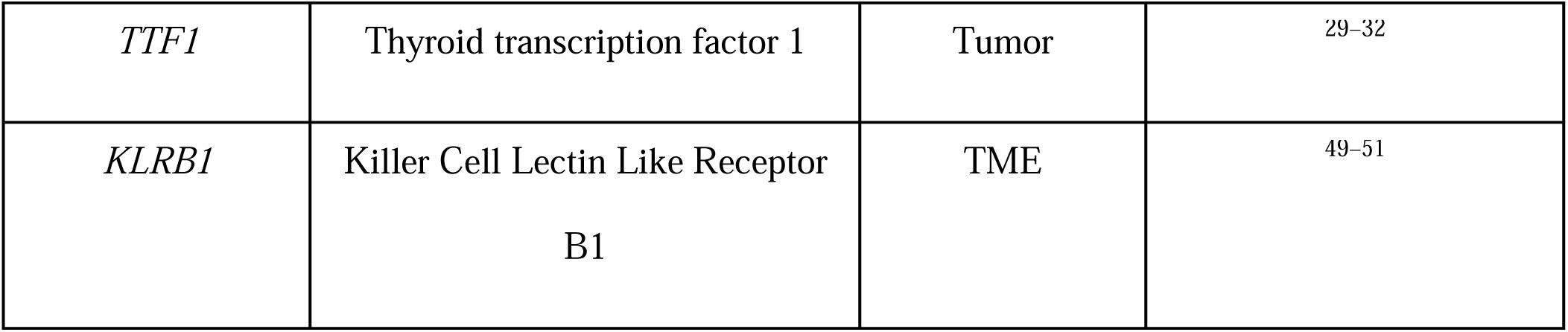
Genes found to be under- or over-expressed in tumor and/or TME regions in LUAD in the literature.

**Table 1b:** Genes found to be under under- or over-expressed in tumor and TME regions in the TCGA-LUAD cohort.

| Genes | Gene category | Mean fpkm | Median fpkm | Min. fpkm | Max. fpkm |
| --- | --- | --- | --- | --- | --- |
| *CDH3* | Tumor | 0.02 | 0.013 | 0 | 0.172 |
| *KRT7* | Tumor | 0.881 | 0.859 | 0.351 | 2.298 |
| *TP53I3* | Tumor | 1.273 | 1.253 | 0.332 | 3.604 |
| *TICAM1* | TME | 1.513 | 1.44 | 0.482 | 3.562 |
| *CXCL13* | TME | 2.401 | 2.276 | 0.797 | 8.245 |
| *CDKN2A* | Tumor | 2.55 | 2.377 | 0.619 | 7.281 |
| *FAM189A1* | Tumor | 4.011 | 3.8 | 2.12 | 9.219 |
| *TTF1* | Tumor | 5.623 | 5.045 | 2.282 | 20.554 |
| *NAPSA* | Tumor | 9.412 | 7.403 | 0.3 | 44.973 |
| *KLRB1* | TME | 11.053 | 5.292 | 0.387 | 113.49 |
| *CD8A* | TME | 16.658 | 14.792 | 4.996 | 53.298 |
| *SLC47A1* | Tumor | 62.297 | 56.398 | 25.308 | 210.073 |

*CDKN2A* ^58–64^, frequently lost in LUAD, drives cell cycle deregulation and poor outcomes, whereas *TP53I3* (PIG3) ^65–69^, a p53-inducible DNA damage response gene, promotes proliferation, invasion, and therapy resistance. *TICAM1* ^70–74^ links Toll-like receptor signaling to type I interferon pathways and has emerged as a prognostic marker, while *CDH3* (P-cadherin) ^75–77^ enhances epithelial–mesenchymal transition and metastatic potential. *SLC47A1* (MATE1) ^78–80^ influences chemotherapeutic sensitivity as a solute transporter, and *FAM189A1*, a multi-pass transmembrane protein, has been reported to be upregulated in LUAD and implicated in treatment adaptation ^81–83^.

### Model Evaluation of Gene Expression

The performance metrics for the predicted expression of 12 LUAD biomarkers from the testing set are shown in Table 2. The model demonstrated strong predictive ability for *NAPSA*, *TP53I3*, and *SLC47A1*, that achieved AUC values of 0.92, 0.84, and 0.84, respectively. All three genes showed high accuracy (0.85), with *TP53I3* and *SLC47A1* achieving consistently strong precision and recall (0.85), indicating confident classification of expression status across samples. *KLRB1* and *FAM189A1* also performed well, with AUCs of 0.83 and 0.8, respectively; *KLRB1* showed balanced precision and recall (0.75-0.76), while *FAM189A1* achieved its highest performance in test accuracy (0.75).

**Table 2:** Performance metrics for the predicted expression of twelve genes in the test set as a function of the best performing kth fold for each gene.

| Biomarker | Best performing fold | AUC CI range | Test error | AUC | Accurac | Precision | Recall | F1 | Gene |
| --- | --- | --- | --- | --- | --- | --- | --- | --- | --- |

|  |  |  |  |  | y |  |  |  | category |
| --- | --- | --- | --- | --- | --- | --- | --- | --- | --- |
| *NAPSA* | 14th | (0.84,1.0) | 0.15 | 0.92 | 0.85 | 0.85 | 0.85 | 0.84 | Tumor |
| *SLC47A1* | 9th | (0.69,1.0) | 0.15 | 0.84 | 0.85 | 0.85 | 0.85 | 0.85 | Tumor |
| *TP53I3* | 14th | (0.69,1.0) | 0.15 | 0.84 | 0.85 | 0.85 | 0.85 | 0.85 | Tumor |
| *KLRB1* | 11th | (0.66,1.0) | 0.25 | 0.83 | 0.75 | 0.76 | 0.75 | 0.75 | TME |
| *FAM189A1* | 12th | (0.61,1.0) | 0.25 | 0.8 | 0.75 | 0.75 | 0.73 | 0.73 | Tumor |
| *TICAM1* | 13th | (0.61,0.96) | 0.25 | 0.78 | 0.75 | 0.76 | 0.73 | 0.73 | TME |
| *CD8A* | 14th | (0.58,0.96) | 0.3 | 0.77 | 0.7 | 0.7 | 0.7 | 0.69 | TME |
| *CXCL13* | 9th | (0.55,0.96) | 0.35 | 0.76 | 0.65 | 0.67 | 0.65 | 0.64 | TME |
| *TTF1* | 12th | (0.47,0.96) | 0.30 | 0.72 | 0.7 | 0.84 | 0.7 | 0.65 | Tumor |
| *CDH3* | 9th | (0.41,0.91) | 0.35 | 0.66 | 0.65 | 0.68 | 0.65 | 0.61 | Tumor |
| *KRT7* | 8th | (0.38,0.92) | 0.35 | 0.65 | 0.65 | 0.68 | 0.65 | 0.62 | Tumor |
| *CDKN2A* | 11th | (0.38,0.89) | 0.35 | 0.64 | 0.65 | 0.65 | 0.63 | 0.63 | Tumor |

Moderate predictive performance was observed for *CD8A* and *TICAM1*, with AUCs of 0.77 and 0.78, respectively. Both genes maintained test accuracies above 0.70, and *TICAM1* showed good precision (∼0.76). *CXCL13* and *TTF1* were predicted with slightly lower AUCs (0.76 and 0.72, respectively), but *TTF1* retained high precision (0.84). Finally, *KRT7*, *CDH3*, and *CDKN2A* had lower AUC values, ranging from 0.64 to 0.66, though all three still achieved test accuracies of 0.65 and demonstrated moderate classification performance across other metrics.

### Visualization and Interpretation of Biomarker Expression

To further investigate the model’s spatial attention and interpretability of ROIs, we examined the attention heatmaps produced by XpressO’s HeatMaps module for test WSIs across biomarkers. Depending on the model’s association of the biomarkers’ expression with the tissue-morphological patterns, a biomarker could be predicted as either high or low-expressed in several WSI-patches, where red-patches indicate regions that influence the model’s prediction the most, followed by yellow and green-patches that have moderate and low influence on that prediction (Figures 1-5 and Supplementary Figures S1-S11). The p_0 and p_1 indicate probabilities for the predicted class (high or low-expression) using top-10 patches and non-predicted class (low or high expression) using bottom-10 patches (Table 3). The heatmaps revealed spatial associations between predicted biomarker expression and histomorphologic features in tumor and TME regions (Figures 1-5 and Supplementary Figures S1-S11). The following subsections illustrate representative examples for individual biomarkers and biomarker pairs, linking predicted expression with corresponding histomorphologic patterns.

**Figure 1.**
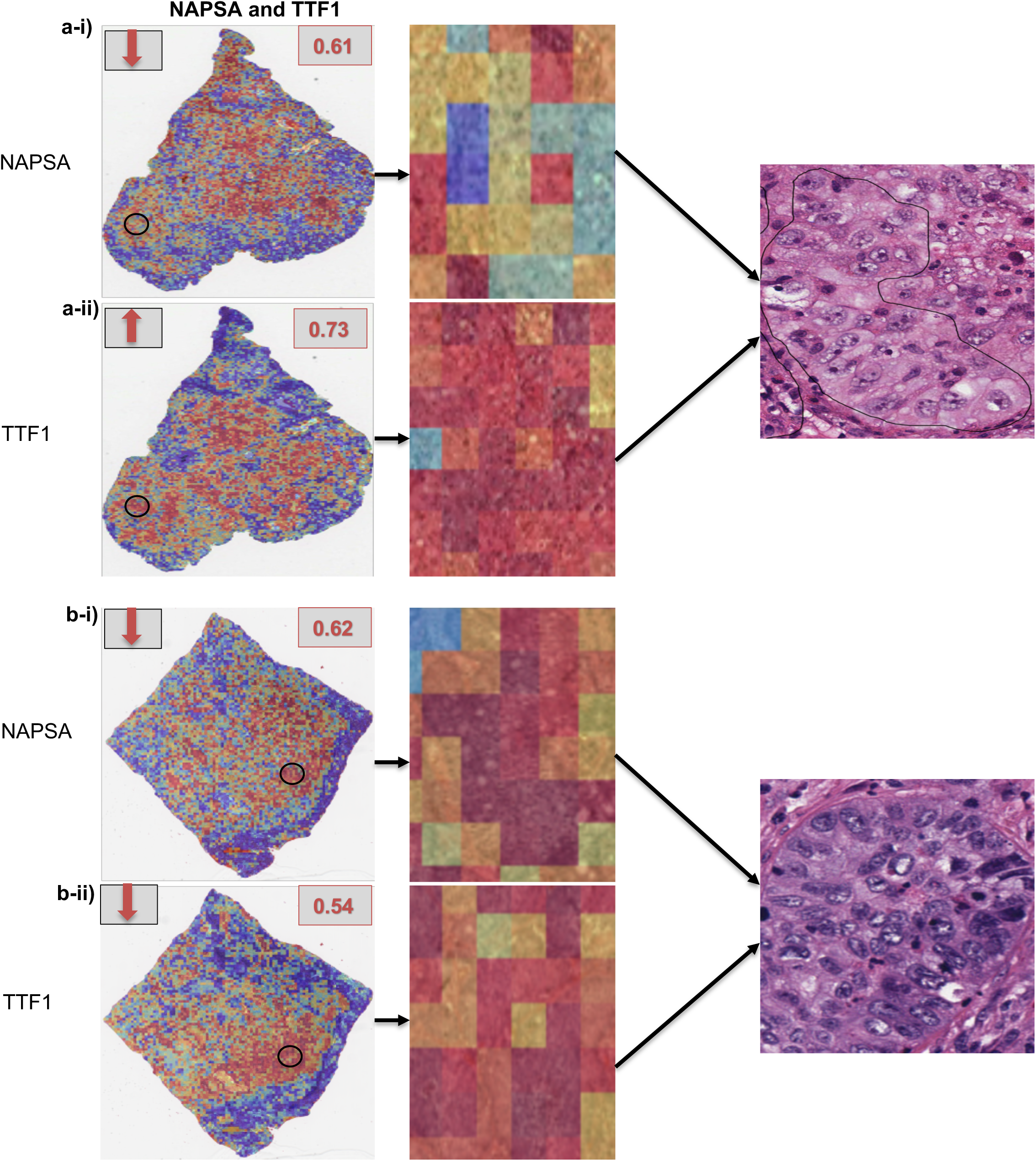
Attention heatmaps and corresponding H&E views for *NAPSA* and *TTF1* expression predictions in LUAD whole-slide images. (a) Case (TCGA-55-A493) showing low *NAPSA* expression (a-i) and high *TTF1* expression (a-ii). (b) Case (TCGA-75-7027 ) showing low expression for both *NAPSA* (b-i) and *TTF1* (b-ii). High or low expression status is indicated by red arrows, with the predicted probability of high or low expression (*p_0*) shown in boxes. Black circles mark regions of interest selected for zoom-in visualization.

**Table 3.** Model-predicted biomarker expression status across TCGA lung cancer samples. For each TCGA case, the model predicts whether the biomarker is in high or low expression state; where pred_0 = Predicted expression class (high/low expression) using the top-10 attended patches; p_0 = Probability of the predicted class using top-10 attention; pred_1 = Predicted expression class (high/low) using the bottom-10 attended patches; p_1 = Probability of the predicted class using bottom-10 attention. Associated figure panels (e.g., 1-a-i, S3-b-ii) point to corresponding heatmaps and H&E visualization.

| TCGA ID | Biomarker | pred_0 | p_0 | Pred_1 | p_1 | Figure |
| --- | --- | --- | --- | --- | --- | --- |
| TCGA-55-A493 | *NAPSA* | low_expression | 0.61 | high_expression | 0.39 | 1-a-i |
|  | *TTF1* | high_expression | 0.73 | low_expression | 0.27 | 1-a-ii |
| TCGA-75-7027 | *NAPSA* | low_expression | 0.62 | high_expression | 0.38 | 1-b-i |
|  | *TTF1* | low_expression | 0.54 | high_expression | 0.46 | 1-b-ii |
| TCGA-55-1592 | *NAPSA* | low_expression | 0.8 | high_expression | 0.2 | S1-a-i |
|  | *TTF1* | high_expression | 0.73 | low_expression | 0.27 | S1-a-ii |
| TCGA-44-5643 | *CD8A* | high_expression | 0.88 | low_expression | 0.17 | S2-a-i |
|  | *KRT7* | high_expression | 0.74 | low_expression | 0.26 | S2-a-ii |
| TCGA-49-AAR3 | *CD8A* | high_expression | 0.61 | low_expression | 0.39 | 2-a-i |
|  | *KRT7* | low_expression | 0.69 | high_expression | 0.31 | 2-a-ii |
| TCGA-69-7973 | *CD8A* | high_expression | 0.82 | low_expression | 0.18 | 2-b-i |
|  | *KRT7* | high_expression | 0.78 | low_expression | 0.22 | 2-b-ii |
| TCGA-44-8119 | *KRT7* | high_expression | 0.53 | low_expression | 0.47 | 3-a-i |
|  | *CDKN2A* | low_expression | 0.76 | high_expression | 0.24 | 3-a-ii |
| TCGA-97-A4M5 | *KRT7* | high_expression | 0.77 | low_expression | 0.23 | 3-b-i |
|  | *CDKN2A* | low_expression | 0.66 | high_expression | 0.34 | 3-b-ii |
| TCGA-55-7907 | *NAPSA* | high_expression | 0.54 | low_expression | 0.46 | 4-b-i |
|  | *CDKN2A* | low_expression | 0.55 | high_expression | 0.45 | 4-b-ii |
| TCGA-55-8506 | *NAPSA* | low_expression | 0.79 | high_expression | 0.21 | S3-a-i |
|  | *CDKN2A* | low_expression | 0.63 | high_expression | 0.37 | S3-a-ii |
| TCGA-97-A4M1 | *NAPSA* | high_expression | 0.91 | low_expression | 0.09 | 4-a-i |
|  | *CDKN2A* | low_expression | 0.75 | high_expression | 0.25 | 4-a-ii |
| TCGA-97-A4M5 | *NAPSA* | high_expression | 0.52 | low_expression | 0.48 | S3-b-i |
|  | *CDKN2A* | low_expression | 0.66 | high_expression | 0.34 | S3-b-ii |
| TCGA-69-7973 | *TTF1* | low_expression | 0.52 | high_expression | 0.48 | 5-a-i |
|  | *KRT7* | high_expression | 0.78 | low_expression | 0.22 | 5-a-ii |
| TCGA-69-8254 | *TTF1* | high_expression | 0.77 | low_expression | 0.23 | S4-a-i |
|  | *KRT7* | high_expression | 0.63 | low_expression | 0.37 | S4-a-ii |
| TCGA-L9-A50W | *TTF1* | low_expression | 0.6 | high_expression | 0.4 | 5-b-i |
|  | *KRT7* | high_expression | 0.89 | low_expression | 0.11 | 5-b-ii |
| TCGA-05-4397 | *TP53I3* | high_expression | 0.87 | low_expression | 0.13 | S5-a |
| TCGA-78-7153 |  | low_expression | 0.91 | high_expression | 0.09 | S5-b |
| TCGA-86-8075 |  | low_expression | 0.56 | high_expression | 0.44 |  |
| TCGA-L9-A443 |  | low_expression | 0.54 | high_expression | 0.46 |  |
| TCGA-67-4679 |  | high_expression | 0.64 | low_expression | 0.36 |  |
| TCGA-L4-A4E6 |  | low_expression | 0.60 | high_expression | 0.40 |  |
| TCGA-97-A4M5 | *TICAM1* | high_expression | 0.86 | low_expression | 0.14 | S6-a |
| TCGA-55-A493 |  | low_expression | 0.52 | high_expression | 0.48 |  |
| TCGA-69-8254 |  | low_expression | 0.57 | high_expression | 0.43 | S6-b |
| TCGA-44-5643 |  | high_expression | 0.82 | low_expression | 0.18 |  |
| TCGA-97-8172 | *SLC47A1* | low_expression | 0.97 | high_expression | 0.03 | S7 |
| TCGA-67-4679 |  | low_expression | 0.98 | high_expression | 0.02 |  |
| TCGA-86-8075 |  | low_expression | 0.98 | high_expression | 0.02 |  |
| TCGA-05-4397 | *FAM189A1* | high_expression | 0.96 | low_expression | 0.04 | S8-a |
| TCGA-75-7027 |  | high_expression | 0.68 | low_expression | 0.32 |  |
| TCGA-97-8179 |  | low_expression | 0.77 | high_expression | 0.23 | S8-b |
| TCGA-75-7027 | *CXCL13* | low_expression | 0.69 | high_expression | 0.31 | S9-a |
| TCGA-78-7153 |  | high_expression | 0.61 | low_expression | 0.39 | S9-b |
| TCGA-55-A493 |  | low_expression | 0.73 | high_expression | 0.27 |  |
| TCGA-97-A4M5 | *KLRB1* | low_expression | 0.51 | high_expression | 0.49 | S10-a |
| TCGA-97-8172 |  | low_expression | 0.60 | high_expression | 0.40 |  |
| TCGA-78-7153 |  | low_expression | 0.72 | high_expression | 0.28 |  |
| TCGA-L4-A4E6 |  | high_expression | 0.69 | low_expression | 0.31 | S10-b |
| TCGA-55-A493 | *CDH3* | high_expression | 0.55 | low_expression | 0.45 | S11-a |
| TCGA-L9-A443 |  | low_expression | 0.63 | high_expression | 0.37 | S11-b |
| TCGA-97-8179 |  | high_expression | 0.66 | low_expression | 0.34 |  |
| TCGA-69-8254 |  | high_expression | 0.58 | low_expression | 0.42 |  |
| TCGA-67-4679 |  | high_expression | 0.50 | low_expression | 0.50 |  |

#### a) NAPSA and TTF1

*NAPSA* (Napsin A) and *TTF1*, two well-established markers of LUAD, were frequently predicted to have expression heterogeneity and exhibited high attention weights within similar tumor regions (Table 3, Fig 1-a,b and Supplementary Fig S1). In WSIs TCGA-55-A493 and TCGA-55-1592, *TTF1* was highly expressed while *NAPSA* showed low expression. In WSI TCGA-75-7027, both markers exhibited low expression. Heatmaps highlighted a focused spatial pattern for *TTF1*, consistent with its stable nuclear localization in the bulk adenocarcinoma clone (Fig 1-a,b, Supplementary Fig S1). In contrast, *NAPSA*’s scattered expression appeared more diffuse within tumor regions, reflecting its cytoplasmic granular staining and possible dilution by adjacent normal pneumocytes (Fig 1-a,b, Supplementary Fig S1) ^84,85^. Morphologically, high-*TTF1* regions showed cohesive gland-forming tumor nests with uniform nuclear features, while low-*NAPSA* areas often corresponded to poorly differentiated foci lacking clear cytoplasmic granularity (Fig 1-a,b, Supplementary Fig S1). These observations are consistent with known histopathologic features and demonstrate the model’s capacity to distinguish subcellular expression patterns directly from WSIs ^86^. While primarily used as diagnostic markers, their heterogeneous expression may also reflect underlying clonal diversity, which has been associated with variable clinical outcomes ^87^.

#### b) CD8A and KRT7

The predicted expression of *CD8A* was consistently high across the three WSIs evaluated (TCGA-44-5643, TCGA-49-AAR3, and TCGA-69-7973) in the test set (Table 3, Fig 2-a,b and Supplementary Fig S2). *KRT7* expression was also predicted to be high in two of these slides (TCGA-44-5643 and TCGA-69-7973) (Table 3, Fig 2a and Supplementary Fig S2), with only one case (TCGA-49-AAR3) (Table 3 and Fig 2b) showing low predicted *KRT7*. Notably, in TCGA-44-5643 and TCGA-69-7973, where both *CD8A* and *KRT7* expression were high, the model-generated heatmaps showed spatial co-localization of *CD8A*-positive regions with *KRT7*-positive epithelial nests. Morphologically, this translated to lymphocytic cuffs and infiltrates directly abutting or penetrating keratin-rich tumor islands, a configuration characteristic of an immune-inflamed TME. This observation is consistent with established histopathologic patterns in LUAD, where a *CD8A* rim penetrating *KRT7*-positive nests is indicative of an immune-inflamed TME ^88^. This immune-inflamed architecture is associated with improved response to immunotherapy and favorable prognosis in LUAD ^89^.

**Figure 2.**
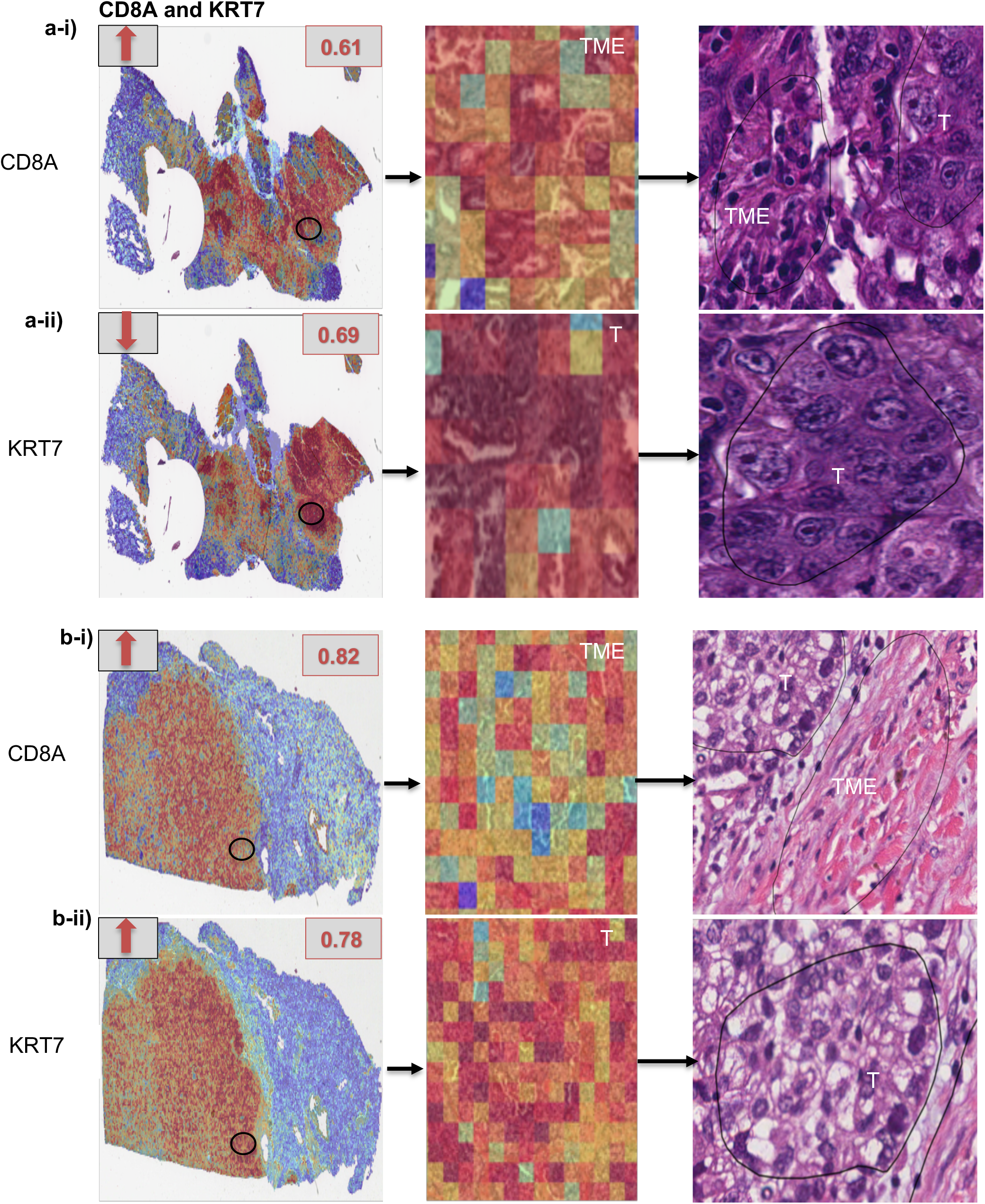
Attention heatmaps and corresponding H&E views for *CD8A* and *KRT7* expression predictions in LUAD whole-slide images. (a) Case (TCGA-49-AAR3) showing high *CD8A* expression (a-i) and low *KRT7* expression (a-ii). (b) Case (TCGA-69-7973) showing high expression for both *CD8A* (b-i) and *KRT7* (b-ii). High or low expression status is indicated by red arrows, with the predicted probability of high or low expression (*p_0*) shown in boxes. Black circles mark regions of interest selected for zoom-in visualization. Areas labeled “T” indicate tumor nests, while areas labeled “TME” denote the surrounding tumor microenvironment.

#### c) KRT7 and CDKN2A

In both test cases analyzed (TCGA-44-8119, TCGA-97-A4M5), *KRT7* expression was predicted to be high, while *CDKN2A* expression was predicted to be low (Table 3 and Figure 3-a,b). This aligns with previously described histologic patterns in lung adenocarcinoma, where regions that are *KRT7*-bright but *CDKN2A*-silent often reflect proliferative LUAD clones with 9p21 loss, a genotype associated with poorer prognosis and potential responsiveness to CDK4/6 or PRMT5-targeted therapies over PD-(L)1 monotherapy ^90,91^. Morphologically, such regions displayed densely cellular epithelial nests with preserved keratin cytoskeleton but minimal intervening stroma, supporting a high proliferative index.

**Figure 3.**
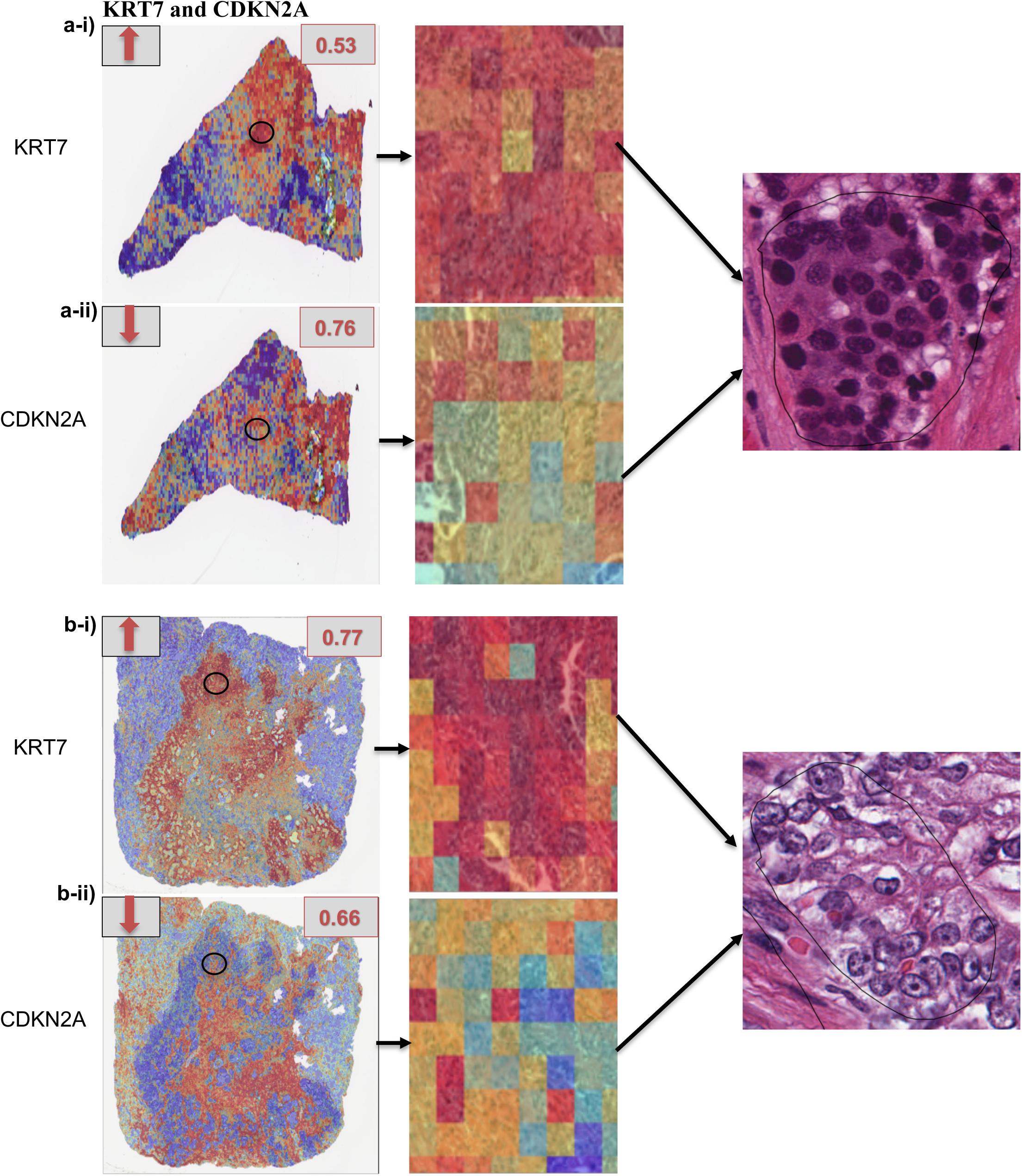
Attention heatmaps and corresponding H&E views for *KRT7* and *CDKN2A* expression predictions in LUAD whole-slide images. (a) Case (TCGA-44-8119) showing high *KRT7* expression (a-i) and low *CDKN2A* expression (a-ii). (b) Case (TCGA-97-A4M5) showing high *KRT7* expression (b-i) and low *CDKN2A* expression (b-ii). High or low expression status is indicated by red arrows, with the predicted probability of high or low expression (*p_0*) shown in boxes. Black circles mark regions of interest selected for zoom-in visualization.

#### d) NAPSA and CDKN2A

Across four test tissues (TCGA-55-7907, TCGA-55-8506, TCGA-97-A4M1, TCGA-97-A4M5), *NAPSA* was predicted as high in three and low in one, while *CDKN2A* was consistently low in each tissue, resulting in the most probable pattern of high *NAPSA* coinciding with low *CDKN2A* expression (Table 3, Fig 4-a,b and Supplementary Fig S3-a,b). This pattern, *NAPSA*–bright yet *CDKN2A*–silent characterizes an alveolar-differentiated but cell-cycle-unleashed LUAD clone, frequently associated with 9p21/MTAP deletions. Morphologically, these regions had abundant, finely vacuolated cytoplasm and centrally placed nuclei, mimicking type II pneumocytes, but arranged in crowded acinar or papillary formations without significant maturation. Tumors with this genotype are associated with poorer prognosis and reduced benefit from PD-(L)1 monotherapy, yet they present actionable vulnerabilities: they are susceptible to CDK4/6 blockade and PRMT5 / MAT2A-directed therapies and are candidates for combination immunotherapy strategies ^90,92,93^.

**Figure 4.**
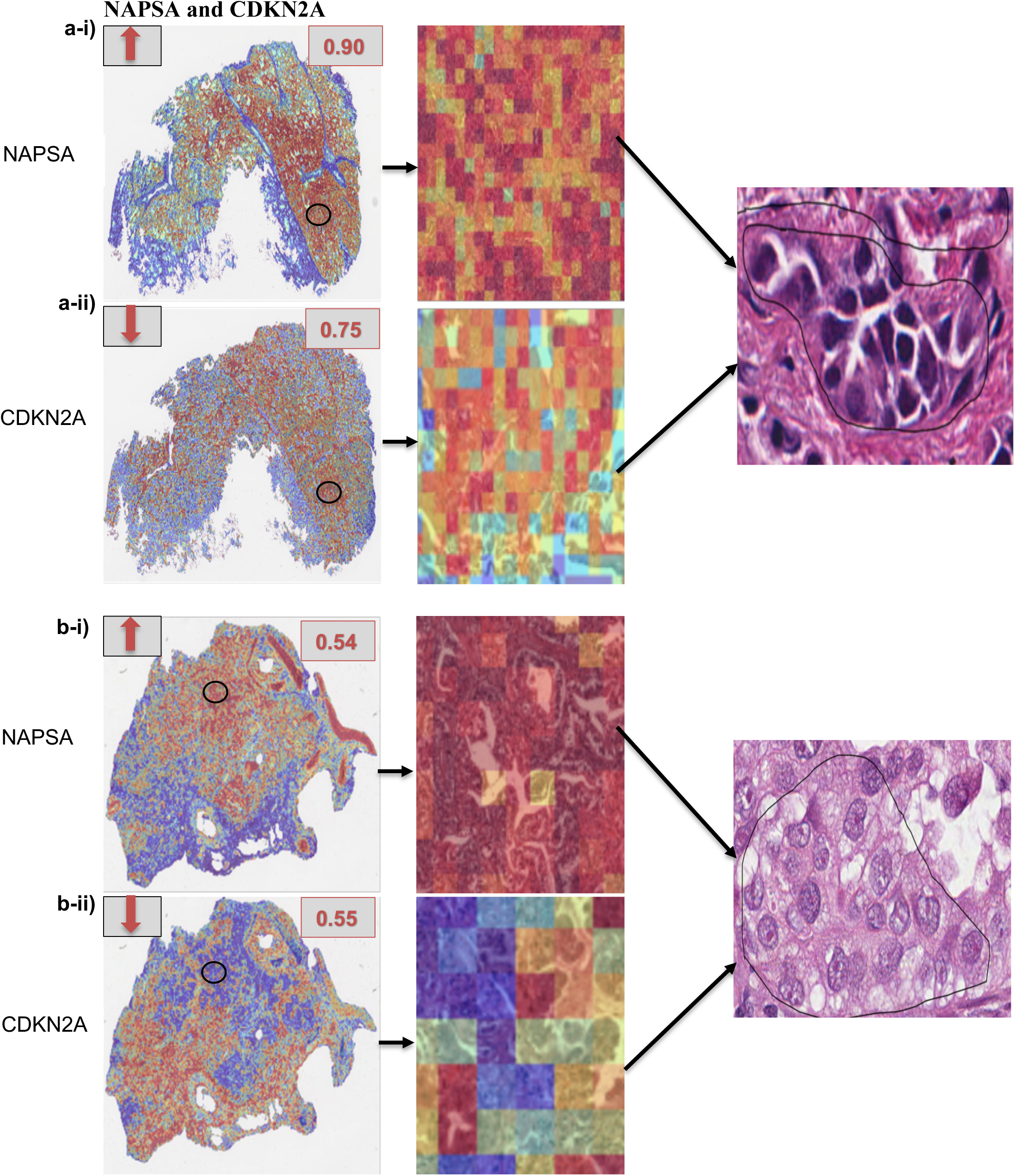
Attention heatmaps and corresponding H&E views for *NAPSA* and *CDKN2A* expression predictions in LUAD whole-slide images. (a) Case (TCGA-97-A4M1) showing high *NAPSA* expression (a-i) and low *CDKN2A* expression (a-ii). (b) Case (TCGA-55-7907) showing high *NAPSA* expression (b-i) and low *CDKN2A* expression (b-ii). High or low expression status is indicated by red arrows, with the predicted probability of high or low expression (*p_0*) shown in boxes. Black circles mark regions of interest selected for zoom-in visualization.

**Figure 5.**
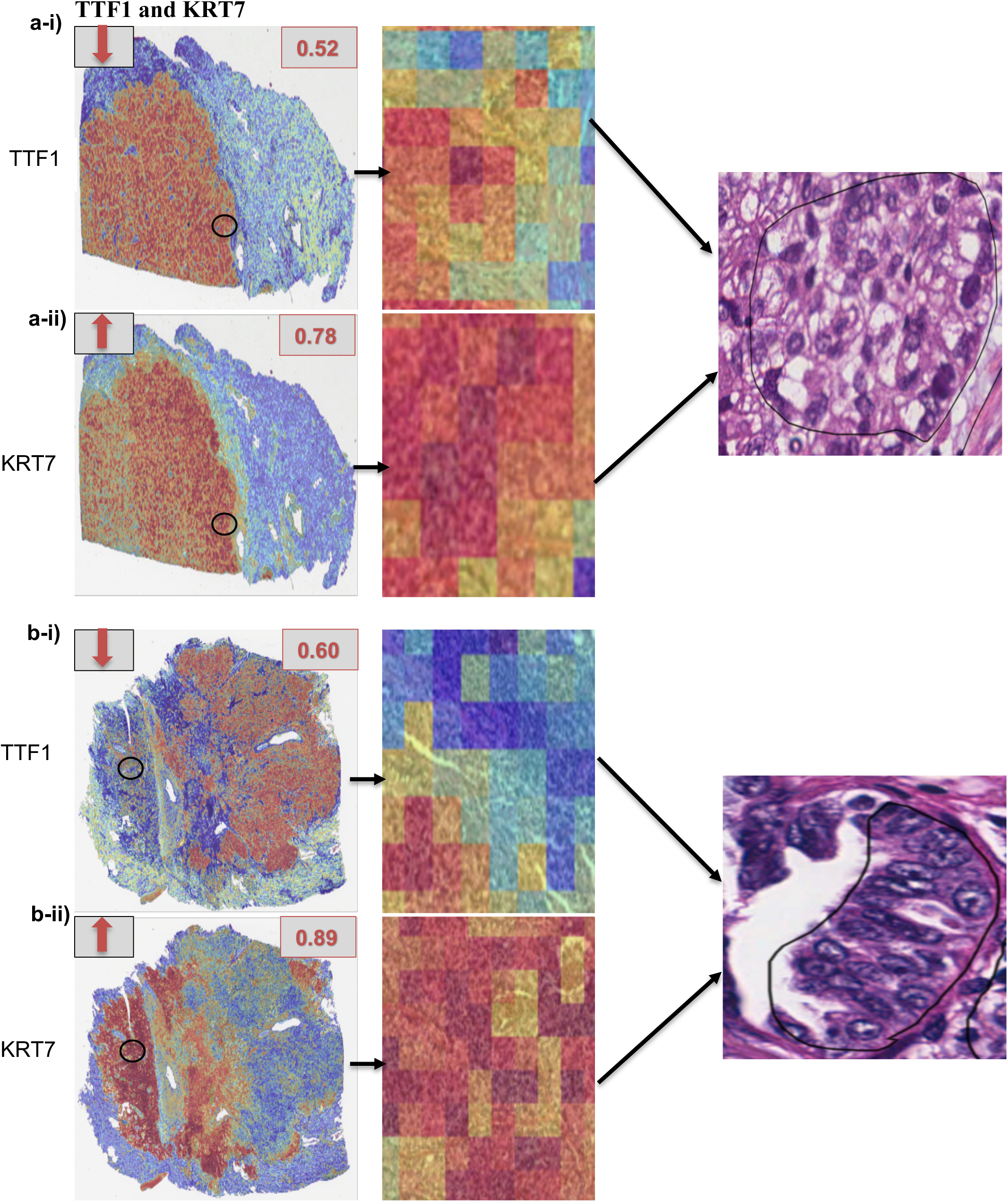
Attention heatmaps and corresponding H&E views for *TTF1* and *KRT7* expression predictions in LUAD whole-slide images. (a) Case (TCGA-69-7973) showing low *TTF1* expression (a-i) and high *KRT7* expression (a-ii). (b) Case (TCGA-L9-A50W) showing low *TTF1* expression (b-i) and high *KRT7* expression (b-ii). High or low expression status is indicated by red arrows, with the predicted probability of high or low expression (*p_0*) shown in boxes. Black circles mark regions of interest selected for zoom-in visualization.

#### e) KRT7 and TTF1

In tissues TCGA-69-7973, TCGA-69-8254, and TCGA-L9-A50W, *KRT7* expression was consistently high, while *TTF1* (NKX2-1) showed variable expression, with low or scattered levels (Table 3, Fig 5-a,b, Supplementary Fig S4). This pattern of sharp *KRT7* alongside reduced or dispersed *TTF1* suggests a histologic landscape containing both normal type II pneumocytes scattered within the tumor and intra-tumoral clones that have selectively downregulated NKX2-1 while retaining keratin cytoskeletal integrity ^94^. Morphologically, this was seen as irregular nests and cords of keratin-rich cells interspersed with alveolar spaces lined by residual *TTF1*-positive pneumocytes.

#### f) TP53I3

*TP53I3*, a transcriptional target of the tumor suppressor TP53, plays a role in oxidative stress response and DNA damage-induced apoptosis ^95^. Predicted expression of *TP53I3* was spatially enriched in areas with architectural disruption or nuclear atypia (Table 3 and Supplementary Fig S5), suggesting that the model may detect regions undergoing genotoxic stress or p53 pathway activation, both of which have implications for tumor aggressiveness and treatment response in LUAD ^96^. Regions with high *TP53I3* expression localized to tumor nests with marked nuclear pleomorphism and structural disruption, consistent with stress-induced p53 signalling (Supplementary Fig S5-a) ^97^, while areas with low *TP53I3* expression showed preserved epithelial organization and uniform nuclei, appearing morphologically uniform and possibly indicative of lower DNA damage response activity (Supplementary Fig S5-b) ^98^.

#### g) TICAM1

*TICAM1*, a signalling adaptor in the TLR3 and cGAS–STING pathways, is increasingly recognized as a marker of innate immune activation in NSCLC ^99,100^. In our analysis, *TICAM1* expression localized to immune-infiltrated tumor borders and stromal pockets (Table 3 and Supplementary Fig S6), consistent with its inclusion in cGAS–STING pathway-associated gene (CSPAG) signatures that stratify NSCLC patients into high- and low-risk groups ^101,102^. Regions with high *TICAM1* expression showed dense lymphocytic cuffs and immune infiltrates at tumor– stromal interfaces, reflecting an immune-“hot” phenotype associated with robust cGAS–STING signaling and favorable outcomes (low-risk group) (Supplementary Fig S6-a)^102^. In contrast, low *TICAM1* expression was observed in morphologically cold tumor nests with limited stromal immune infiltration, consistent with diminished innate immune activation and poorer prognosis (high-risk group) (Supplementary Fig S6-b) ^102^. These findings reinforce *TICAM1*’s role as a classifier of tumor immunogenicity and support its emerging relevance as both a prognostic marker and a therapeutic target in LUAD ^70^.

#### h) SLC47A1

*SLC47A1*, a solute carrier transporter and actionable therapeutic target, has been shown to sensitize cancer cells to platinum–acridine agents, a novel class of chemotherapeutics with enhanced DNA-damaging potential ^103^. In our model, predicted low expression of *SLC47A1* was consistent across three LUAD test cases (Table 3 and Supplementary Fig. S7). Attention heatmaps localized these predictions to well-differentiated glandular regions with preserved acinar structures and uniform nuclei, indicating that reduced *SLC47A1* expression can occur within histologically stable epithelium. This highlights an additional layer of expression-level heterogeneity in LUAD that may be relevant for therapeutic response ^104^. Low SLC47A1 expression has also been associated with resistance to conventional chemotherapy and poorer survival outcomes ^105^.

#### i) FAM189A1

*FAM189A1*, a CD20-like multi-pass transmembrane protein, has been implicated in both lung and hematologic malignancies through mechanisms such as gene-disrupting translocations and promoter methylation, suggesting a role in cell signaling and tumor adaptation ^82,83^. In LUAD, it has also been reported as upregulated in radioresistant NSCLC sublines ^83^. In our analysis, regions with high predicted *FAM189A1* expression localized to cohesive, gland-forming acinar nests with preserved epithelial organization, consistent with a role in maintaining signaling integrity and epithelial polarity ^82^ (Table 3 and Supplementary Fig S8-a). By contrast, regions with low predicted expression were enriched in papillary structures characterized by elongated nuclei, nuclear crowding, and irregular alignment along fibrovascular cores, suggesting a shift toward less cohesive growth and altered epithelial architecture ^82^ (Table 3 and Supplementary Fig S8-b). Together, these findings highlight *FAM189A1* as a potentially important biomarker of tumor plasticity and therapeutic resistance in LUAD ^83^.

#### j) CXCL13

*CXCL13*, a B cell–attracting chemokine, plays a critical role in tertiary lymphoid structure (TLS) formation and has been associated with improved immunotherapy response in NSCLC ^52^. In our analysis, predicted expression of *CXCL13* was found localized to stromal patches adjacent to tumor epithelial fronts, frequently overlapping with lymphoid aggregates or immune-infiltrated regions (Table 3 and Supplementary Fig S9). Morphologically, high-expression regions showed densely clustered lymphoid aggregates within stroma, consistent with TLS-like organization (Supplementary Fig S9-b), whereas low-expression cases displayed more dispersed stromal lymphocytes without clear clustering (Supplementary Fig S9-a) ^52^. These findings align with reports that *CXCL13*-expressing T and B cells are enriched in TLS-positive tumors and highlight the model’s sensitivity to spatially organized immune niches ^106^. Additionally, high CXCL13 expression and TLS presence have been associated with enhanced immunotherapy response and improved prognosis in LUAD ^107^.

#### k) KLRB1

*KLRB1*, which encodes CD161, is a marker of natural killer (NK) cells and subsets of memory-like CD8 and CD4 T cells, particularly tissue-resident T cells implicated in antitumor immunity ^108^. In LUAD, *KLRB1* expression has been associated with favorable prognosis and enhanced immune infiltration. In our cohort, three test cases were predicted to have low *KLRB1* expression, with attention maps predominantly highlighting immune-poor stroma and tumor epithelial regions lacking dense lymphocytic infiltrates (Table 3 and Supplementary Fig S10-a). These areas were characterized by sparse or absent small, round lymphocyte nuclei within the stromal matrix. In contrast, the single high-expression case showed model attention concentrated in lymphocyte-rich stromal zones and peritumoral lymphoid aggregates, consistent with organized immune niches (Table 3 and Supplementary Fig S10-a) . These morphologic features align with prior reports linking reduced *KLRB1* activity to diminished immune cell presence, while higher expression in the literature is associated with enriched lymphocyte-rich microenvironments and improved response to checkpoint blockade in NSCLC ^49^.

#### l) CDH3

*CDH3* (P-cadherin), a calcium-dependent cell–cell adhesion protein typically expressed in basal or progenitor epithelial cells, is frequently upregulated in lung adenocarcinoma (LUAD) and has been associated with increased tumor cell proliferation, migration, and poor prognosis ^109^. Mechanistically, *CDH3* overexpression can promote tumor aggressiveness by modulating adherens junction stability and activating downstream Wnt/β-catenin signaling pathways ^110^. In our attention heatmap analysis, regions predicted to have high *CDH3* expression corresponded to cohesive nests of malignant epithelial cells with preserved gland-forming architecture, a pattern consistent with its role in maintaining cell–cell adhesion while supporting proliferative tumor growth (Table 3 and Supplementary Fig S11-a) ^111^. Low *CDH3* expression localized to disorganized tumor regions with looser epithelial cohesion, irregular cell borders, and nuclear variability, reflecting weakened junctional integrity (Table 3 and Supplementary Fig S11-b).

Together, these patterns align with *CDH3*’s known role in regulating epithelial adhesion and tumor morphology in LUAD ^109^.

The biomarker pairs identified through our model also align with disease associations and pathways highlighted in STRING (version 12.5) ^112^, reinforcing their biological relevance. For instance, *CD8A*, *CDKN2A*, and *KRT7* showed strong associations with non-small cell lung carcinoma, vulvar carcinoma, and melanoma, with a very low false discovery rate in the Disease-gene Associations (DISEASES) ^113^ (Supplementary Fig. S12a). In this context, the FDR represents the probability that a reported association is a false positive after adjusting for multiple hypothesis testing, and values near zero indicate highly confident enrichment. Literature linked to these markers underscores their translational importance (Supplementary Fig. S12b): case reports highlight how immune contexture and PD-L1–driven dynamics in lung cancer can shape outcomes, implicating *CD8A*, *CDKN2A*, and *KRT7* in pathways that govern both therapeutic resistance and progression across subtypes ^114^. Other studies emphasize their roles in intratumoral heterogeneity that drives drug resistance ^115^ and spatial heterogeneity influencing survival in lung adenocarcinoma patients ^116^. Furthermore, guidelines for diagnostic immunohistochemistry in lung cancer recommend markers like *KRT7* and *CDKN2A* as lineage classifiers, directly supporting their inclusion in predictive modelling ^117^.

Another STRING-derived network connecting *CXCL13*, *KLRB1*, and *CD8A* captures immune regulatory processes with direct relevance to the TME (Supplementary Fig. S13). These markers appear in prognostic gene signatures such as necroptosis-associated immune profiles in hepatocellular carcinoma ^118^, and in studies distinguishing intratumoral vs. circulating lymphoid populations as predictive biomarkers in lung cancer patients undergoing immune checkpoint blockade ^119^. Additional work in single-cell settings reveals their role in defining exhausted CD8+ T cells recognizing tumor neoantigens, consistent with their function as immune infiltration markers in NSCLC ^120^. The inclusion of *KLRB1*, a regulator of NK and T cell function, situates this network at the interface of adaptive and innate immunity, providing a mechanistic complement to our model’s predictions.

Finally, STRING associations among *TTF1*, *KRT7*, *CDKN2A*, and *NAPSA* connect these canonical lung adenocarcinoma markers to both disease associations (NSCLC, intestinal cancer, vulvar carcinoma) and specific pathways such as surfactant metabolism, in which *TTF1* and *NAPSA* play key regulatory roles (Supplementary Fig. S14a). Here again, FDR-adjusted enrichment values were very low, indicating robust statistical confidence. Literature linked to this cluster highlights their clinical importance in molecularly complex or therapy-resistant settings (Supplementary Fig. S14b): ALK-rearranged lung tumors lacking typical lineage markers ^121^, ALK fusion–driven resistance to osimertinib ^122^, and histomorphological transformation from NSCLC to small cell carcinoma after immunotherapy ^123^ all emphasize the diagnostic and prognostic value of these markers. Meta-analyses distinguishing adenocarcinoma from squamous carcinoma further reinforce the utility of *KRT7* and *TTF1* as diagnostic anchors ^124^ . Notably, STRING highlights a local cluster of *NAPSA* and *KRT7* with signet-ring cell adenocarcinoma, a rarer subtype, while in our LUAD cohort, these markers instead aligned with classical adenocarcinoma morphologies, reinforcing their broader lineage-defining role. (Supplementary Fig. S14a). These STRING-derived connections illustrate how the biomarker pairs identified by our model intersect with established cancer pathways, disease associations, and prior studies.

## Discussion

In this study, we present XpressO-Lung, an explainable deep learning model that predicts gene expression heterogeneity spatially in LUAD tumors and their microenvironment directly from H&E-stained Dx-WSIs using bulk-transcriptomics data. The model captured histomorphologic patterns predictive of both immune and epithelial gene expression signatures within LUAD tissues. Our results demonstrate that the model not only recapitulates known LUAD biomarkers, i.e., *NAPSA*, *TTF1*, *KRT7*, *CDKN2A*, *CD8A*, *SLC47A1*, *TP53I3*, *KLRB1*, *FAM189A1*, *TICAM1*, *CXCL13*, *CDH3* but also reveals their biologically meaningful spatial expression patterns either in a paired or individual format, mirroring underlying tumor and TME interactions that can inform patient prognosis. Across these biomarkers, the model’s predictive performance ranged from AUC 0.64-0.92, providing a strong foundation for interpreting the spatial patterns it uncovered across distinct epithelial lineages, proliferative states, and immune–stromal niches in LUAD.

XpressO-Lung uncovered diverse epithelial lineages and proliferative states within LUAD. NAPSA and TTF1 frequently co-localized in cohesive glandular nests, anchoring well-differentiated adenocarcinoma clones ^84–86^, while KRT7 often appeared in sharp keratin-rich nests alongside scattered TTF1, suggesting lineage drift ^93^. In contrast, KRT7-bright/CDKN2A-silent regions marked 9p21-loss proliferative clones ^89–90^, and NAPSA-bright/CDKN2A-silent zones reflected alveolar-differentiated yet cell-cycle-unrestrained phenotypes, both linked to poor prognosis yet potentially targetable by CDK4/6 or PRMT5 inhibitors, highlighting their relevance as prognostic and therapeutic biomarkers ^91–92^. CDH3 was enriched in cohesive nests with intact architecture, whereas its loss accompanied disorganized epithelial growth and has been linked to poorer clinical outcomes ^108^. These spatial phenotypes align with STRING associations connecting NAPSA, TTF1, KRT7, CDKN2A, and CDH3 to surfactant metabolism, cell-cycle regulation, and NSCLC pathways ^118–121^.

Beyond epithelial compartments, the model also mapped immune and stromal microenvironments with clear spatial resolution, offering insights into tumor-immune interactions that modulate patient prognosis. CD8A frequently rimmed KRT7-positive tumor islands, forming lymphocytic cuffs characteristic of an immune-inflamed TME, often linked to better survival ^88^. CXCL13 clustered at peritumoral lymphoid aggregates, consistent with TLS-like niches and improved prognosis^106–107^, while KLRB1 marked lymphocyte-rich stromal pockets and was absent from immune-poor zones ^81^. TICAM1 concentrated at tumor-stromal borders with dense lymphocytic infiltrates, suggesting cGAS–STING activation, whereas its loss delineated immune-cold nests often associated with poor clinical outcomes ^101–102^. These findings mirror STRING-derived networks linking CD8A, CXCL13, and KLRB1 to immune regulatory pathways and checkpoint blockade response, underscoring their value as spatial markers of immune activity ^115–117^.

Finally, XpressO-Lung identified stress-responsive and therapy-relevant niches within LUAD tissue. TP53I3 marked disordered regions with nuclear pleomorphism, hinting at p53 pathway activation and oxidative stress, while its low expression was observed in morphologically uniform zones ^97^. SLC47A1 was uniformly low in well-differentiated acini, echoing its reported role in sensitizing tumors to platinum–acridine therapeutics ^102^. FAM189A1 showed a striking spatial dichotomy, high in cohesive acinar nests and low in papillary structures with disordered polarity consistent with its reported links to therapy adaptation and tumor plasticity ^53^. Together, these patterns illustrate how XpressO-Lung characterizes the spatial architecture of the LUAD tumor–TME interface, revealing how morphology encodes both lineage state and clinically-relevant phenotypes.

While XpressO-Lung demonstrates strong predictive performance and spatial interpretability, certain limitations remain. Although the model is trained to associate global expression with local histologic features, validation at the spatial level remains a challenge. The TCGA-LUAD dataset does not include spatial transcriptomics or immunohistochemistry (IHC) data for most biomarkers analyzed, limiting our ability to directly validate patch-level expression predictions. In future work, we plan to close this gap by performing targeted spatial transcriptomics and multiplexed IHC staining on in-house LUAD tissue cohorts. This will enable finer-grained validation of XpressO-Lung’s attention maps and expression predictions, further strengthening its clinical relevance and increasing its utility for prognostic applications in translational settings.

XpressO-Lung advances the field by offering an accessible, scalable, and interpretable tool for morpho-genomic profiling in LUAD. It bridges histopathology and bulk-transcriptomics using standard clinical slides, circumventing the need for expensive molecular assays, and enabling prognostic biomarker discovery, stratification of immune versus epithelial tumor states, and improved patient-specific therapy selection. In deep learning for computational pathology, interpreting model outputs is as critical as achieving high predictive performance, and XpressO-Lung delivers on both fronts. The model’s framework reliably infers spatial gene expression patterns from histologic images and highlights biologically meaningful tumor-TME interactions that are predictive of patient prognosis. The model was trained, validated, and tested using a carefully structured design that ensured sufficient degrees of freedom for each gene, and incorporated a custom script for gene expression analysis to systematically reveal morphology-expression relationships. By combining rigorous methodology with interpretable outputs, XpressO-Lung offers a powerful complement to molecular profiling, with the potential to support prognostic assessment, inform clinical decision-making, and advance precision medicine in LUAD, especially in low-resource settings.

## Data Availability Statement

All data analyzed during this study are either included in this article or at the repository: https://github.com/skr1/XpressO.

## Acknowledgements

Author Contributions Statement:

Conceptualization SS, XL; Data curation VR, SS, AW, XL; Formal analysis VR, SS, AW, XL; Funding acquisition SS; Investigation SS, VR, XL, LL, AW; Methodology VR, SS, AW, XL; Project administration SS; Resources SS; Software SS, VR; Supervision SS, XL; Validation; Visualization SS, VR, XL, AW; Writing – original draft VR, SS; Writing – review & editing VR, SS, XL, LL, AW

## SUPPLEMENTARY FIGURE LEGENDS

**Supplementary Figure S1. Attention heatmaps and corresponding H&E views for *NAPSA* and *TTF1* expression predictions in LUAD whole-slide images.**

(a) Case (TCGA-55-1592) showing low *NAPSA* expression (a-i) and high *TTF1* expression (a-ii). High or low expression status is indicated by red arrows, with the predicted probability of high or low expression (*p_0*) shown in boxes. Black circles mark regions of interest selected for zoom-in visualization.

**Supplementary Figure S2. Attention heatmaps and corresponding H&E views for *CD8A* and *KRT7* expression predictions in LUAD whole-slide images.**

(a) Case (TCGA-44-5643) showing high *CD8A* expression (a-i) and high *KRT7* expression (a-ii). High or low expression status is indicated by red arrows, with the predicted probability of high or low expression (*p_0*) shown in boxes. Black circles mark regions of interest selected for zoom-in visualization. Areas labeled “T” indicate tumor nests, while areas labeled “TME” denote the surrounding tumor microenvironment.

**Supplementary Figure S3. Attention heatmaps and corresponding H&E views for *NAPSA* and *CDKN2A* expression predictions in LUAD whole-slide images.**

(a) Case (TCGA-55-8506) showing low *NAPSA* expression (a-i) and low *CDKN2A* expression (a-ii). (b) Case (TCGA-97-A4M5) showing high *NAPSA* expression (b-i) and low *CDKN2A* expression (b-ii). High or low expression status is indicated by red arrows, with the predicted probability of high or low expression (*p_0*) shown in boxes. Black circles mark regions of interest selected for zoom-in visualization.

**Supplementary Figure S4. Attention heatmaps and corresponding H&E views for *TTF1* and *KRT7* expression predictions in LUAD whole-slide images.**

(a) Case (TCGA-69-8254) showing high *TTF1* expression (a-i) and high *KRT7* expression (a-ii). High or low expression status is indicated by red arrows, with the predicted probability of high or low expression (*p_0*) shown in boxes. Black circles mark regions of interest selected for zoom-in visualization.

**Supplementary Figure S5. Attention heatmaps and corresponding H&E views for *TP53I3* expression predictions in LUAD whole-slide images.**

(a) Case (TCGA-05-4397) showing high *TP53I3* expression. (b) Case (TCGA-78-7153) showing low *TP53I3* expression High or low expression status is indicated by red arrows, with the predicted probability of high or low expression (*p_0*) shown in boxes. Black circles mark regions of interest selected for zoom-in visualization.

**Supplementary Figure S6. Attention heatmaps and corresponding H&E views for *TICAM1* expression predictions in LUAD whole-slide images.**

(a) Case (TCGA-97-A4M5) showing high *TICAM1* expression. (b) Case (TCGA-69-8254) showing low *TICAM1* expression. High or low expression status is indicated by red arrows, with the predicted probability of high or low expression (*p_0*) shown in boxes. Black circles mark regions of interest selected for zoom-in visualization. Areas labeled “T” indicate tumor nests, while areas labeled “TME” denote the surrounding tumor microenvironment

**Supplementary Figure S7. Attention heatmaps and corresponding H&E views for *SLC47A1* expression predictions in LUAD whole-slide images.**

Case (TCGA-97-8172) showing low *SLC47A1* expression. High or low expression status is indicated by red arrows, with the predicted probability of high or low expression (*p_0*) shown in boxes. Black circles mark regions of interest selected for zoom-in visualization.

**Supplementary Figure S8. Attention heatmaps and corresponding H&E views for *FAM189A1* expression predictions in LUAD whole-slide images.**

(a) Case (TCGA-05-4397) showing high *FAM189A1* expression. (b) Case (TCGA-97-8179) showing low *FAM189A1* expression High or low expression status is indicated by red arrows, with the predicted probability of high or low expression (*p_0*) shown in boxes. Black circles mark regions of interest selected for zoom-in visualization. Areas labeled “T” indicate tumor nests, while areas labeled “TME” denote the surrounding tumor microenvironment.

**Supplementary Figure S9. Attention heatmaps and corresponding H&E views for *CXCL13* expression predictions in LUAD whole-slide images.**

(a) Case (TCGA-75-7027) showing low *CXCL13* expression. (b) Case (TCGA-78-7153) showing high *CXCL13* expression. High or low expression status is indicated by red arrows, with the predicted probability of high or low expression (*p_0*) shown in boxes. Black circles mark regions of interest selected for zoom-in visualization. Areas labeled “T” indicate tumor nests, while areas labeled “TME” denote the surrounding tumor microenvironment.

**Supplementary Figure S10. Attention heatmaps and corresponding H&E views for *KLRB1* expression predictions in LUAD whole-slide images.**

(a) Case (TCGA-97-A4M5) showing low *KLRB1* expression. (b) Case (TCGA-L4-A4E6) showing high *KLRB1* expression. High or low expression status is indicated by red arrows, with the predicted probability of high or low expression (*p_0*) shown in boxes. Black circles mark regions of interest selected for zoom-in visualization. Areas labeled “T” indicate tumor nests, while areas labeled “TME” denote the surrounding tumor microenvironment.

**Supplementary Figure S11. Attention heatmaps and corresponding H&E views for *CDH3* expression predictions in LUAD whole-slide images.**

(a) Case (TCGA-55-A493) showing high *CDH3* expression. (b) Case (TCGA-L9-A443) showing low *CDH3* expression. High or low expression status is indicated by red arrows, with the predicted probability of high or low expression (*p_0*) shown in boxes. Black circles mark regions of interest selected for zoom-in visualization.

**Supplementary Figure S12.** (a) STRING disease–gene associations for *CD8A*, *CDKN2A*, and *KRT7* from DISEASES. (b) Reference publications linked to these markers. Node colors reflect supporting evidence categories; edges (green line) represent text-mining associations.

**Supplementary Figure S13.** STRING network of *CXCL13*, *KLRB1*, and *CD8A* with reference publications. Node colors reflect supporting evidence categories; edges (green line) represent text-mining associations.

**Supplementary Figure S14.** (a) STRING Reactome pathway, network cluster, and disease gene associations for *TTF1*, *KRT7*, *CDKN2A*, and *NAPSA*. (b) Reference publications linked to this cluster. Node colors reflect supporting evidence categories; edges (green line) represent text-mining associations.

