## Supplementary Figures for "Interpretable Deep Learning Reveals Biologically Relevant Spatial Gene Expression Patterns in Lung Tumors and their Microenvironment"

**NAPSA and TTF1**

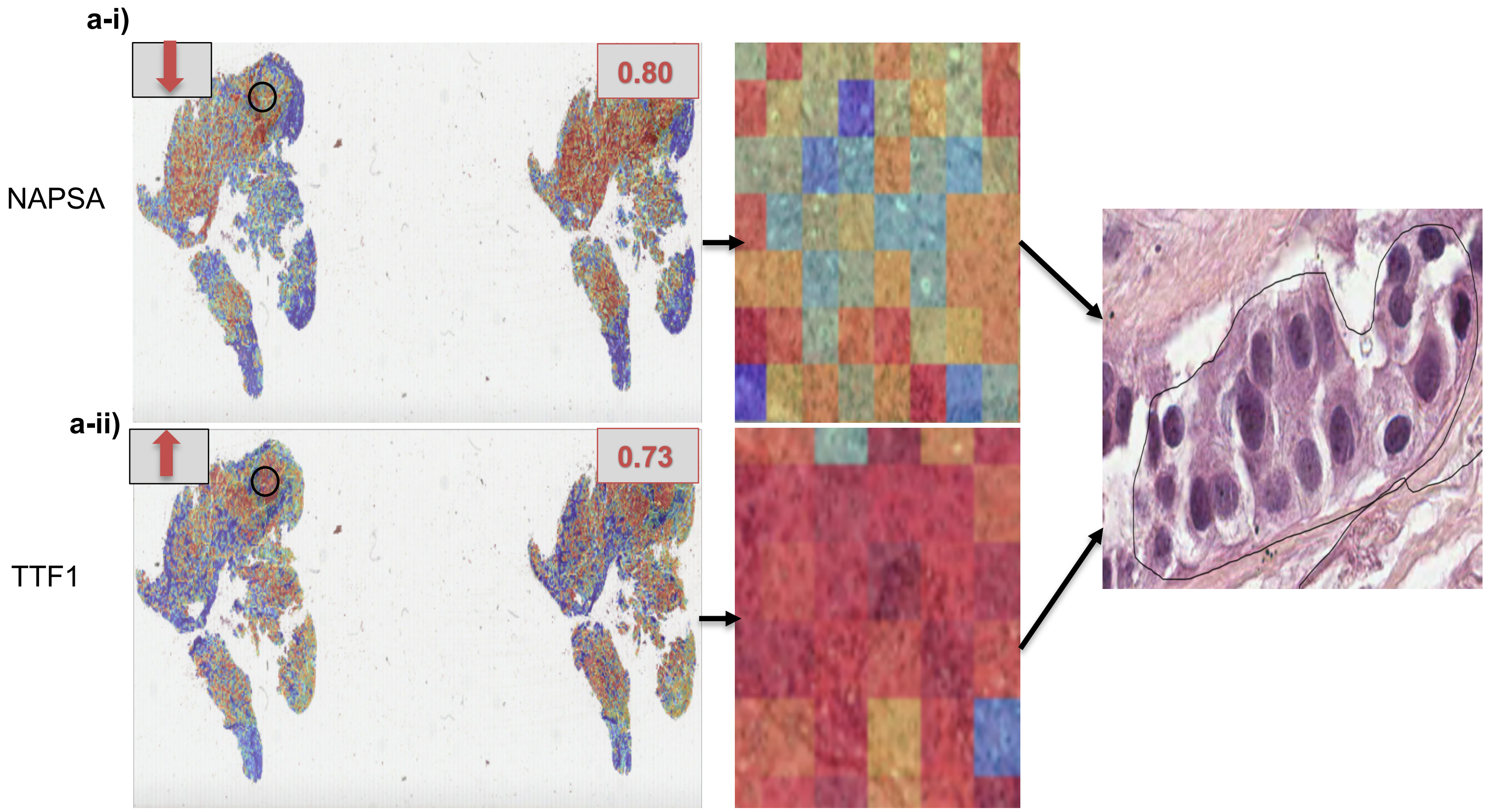


**Supplementary Figure S1**

**CD8A and KRT7**

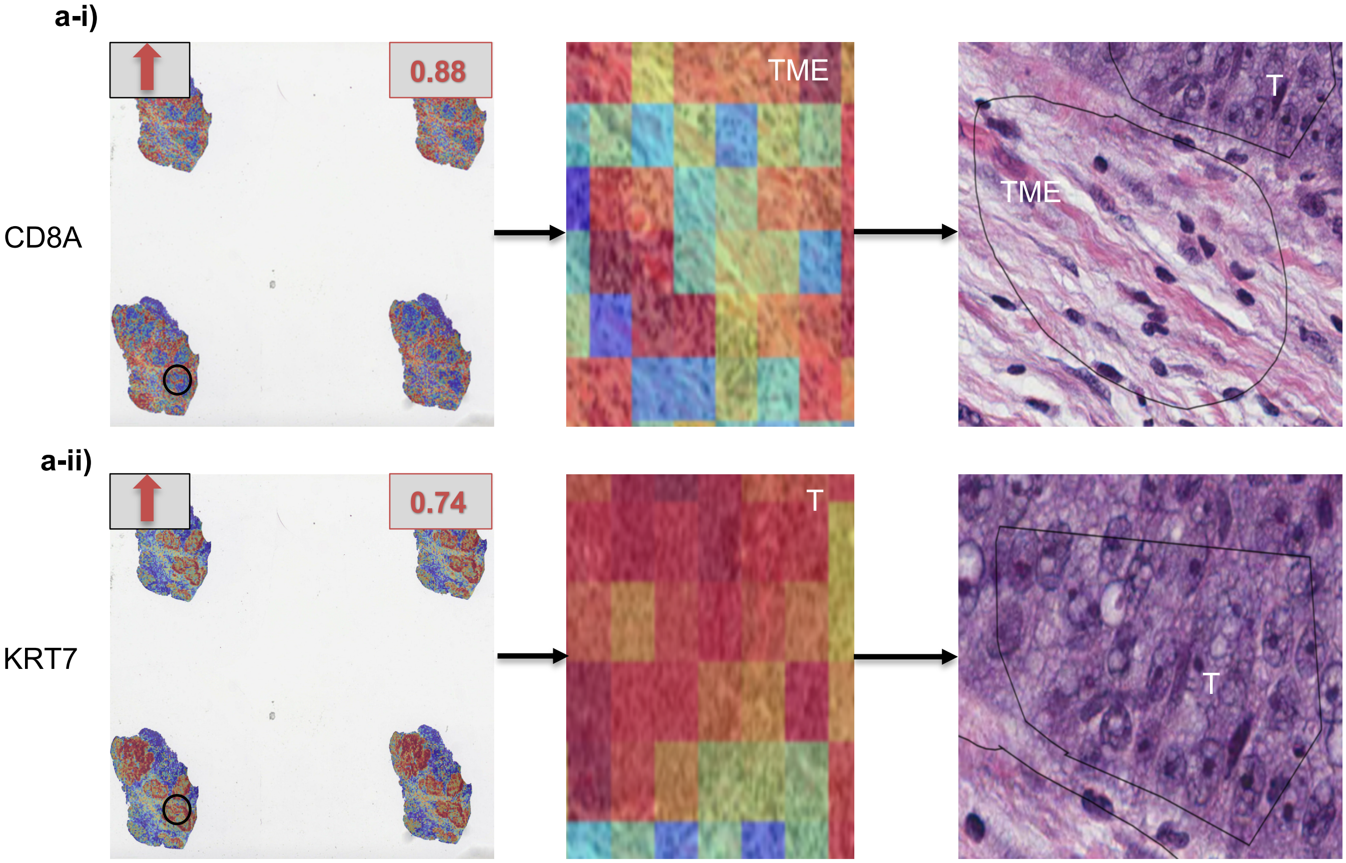


**Supplementary Figure S2**

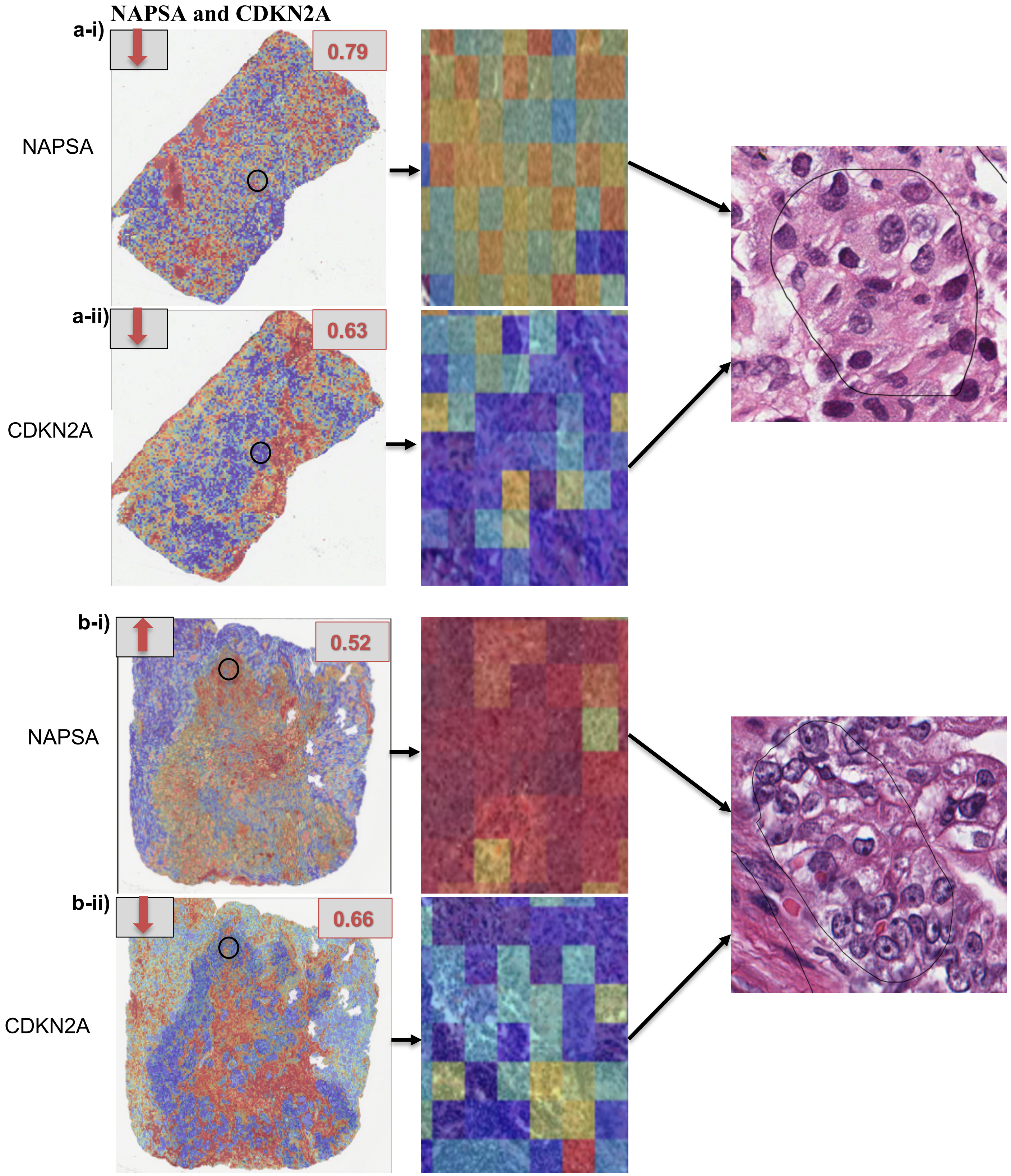


**Supplementary Figure S3**

**TTF1 and KRT7**

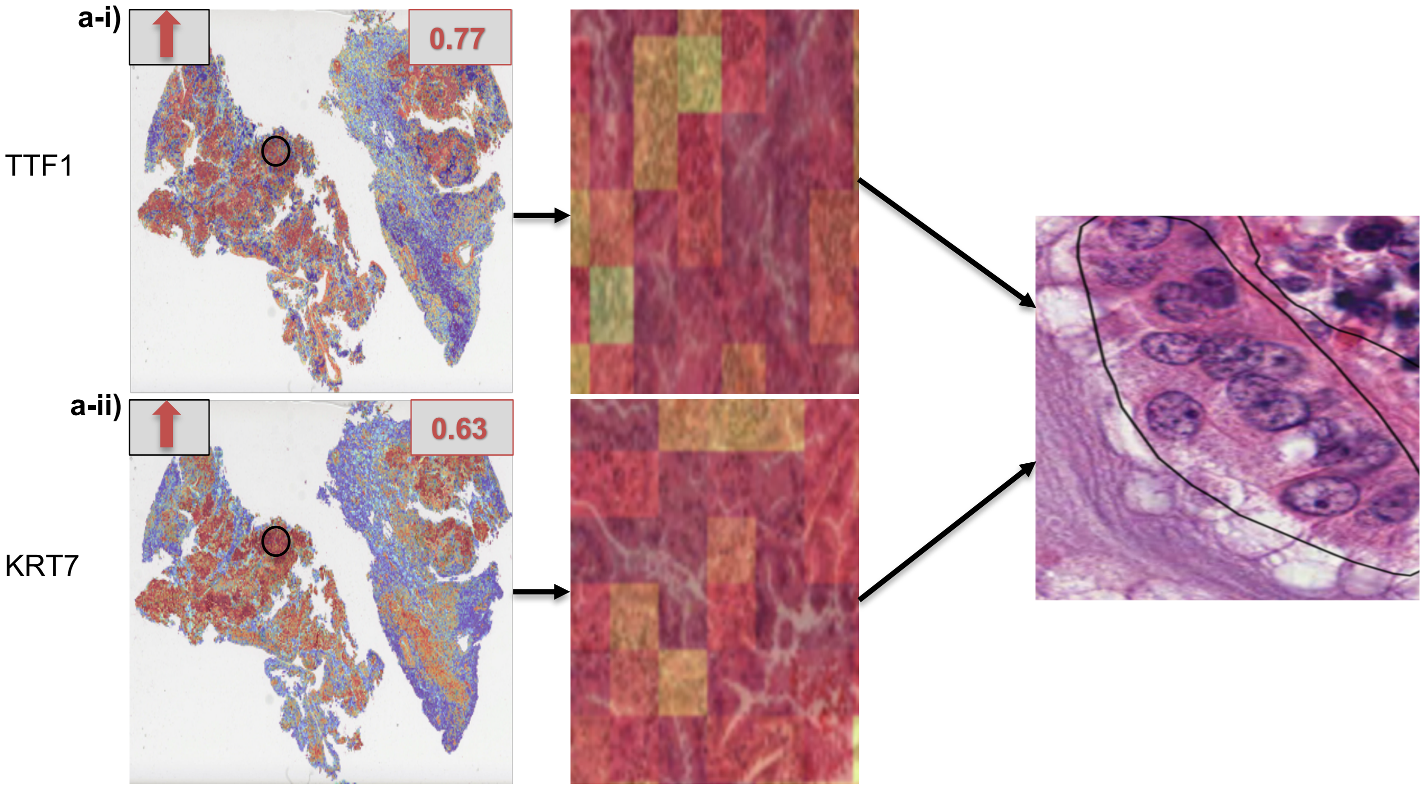


**Supplementary Figure S4**

**TP53I3**

**a)**

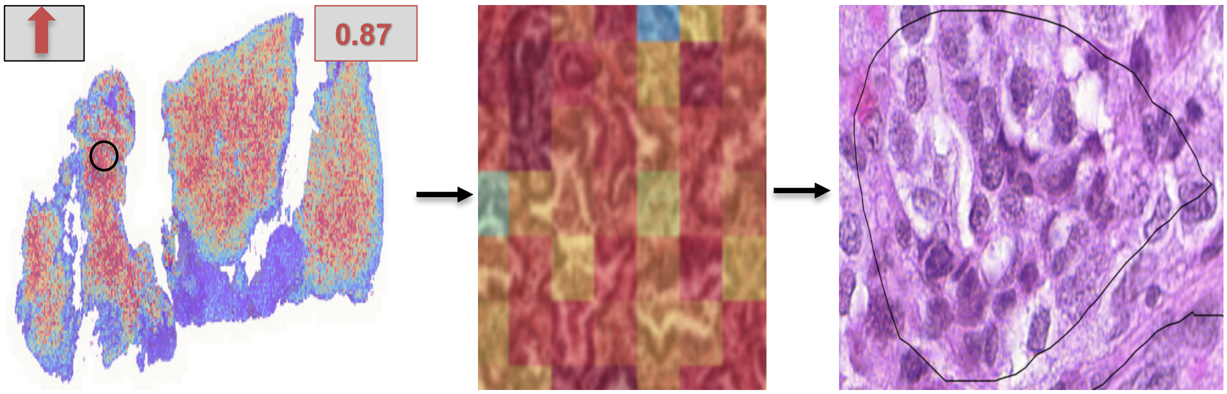


**b)**

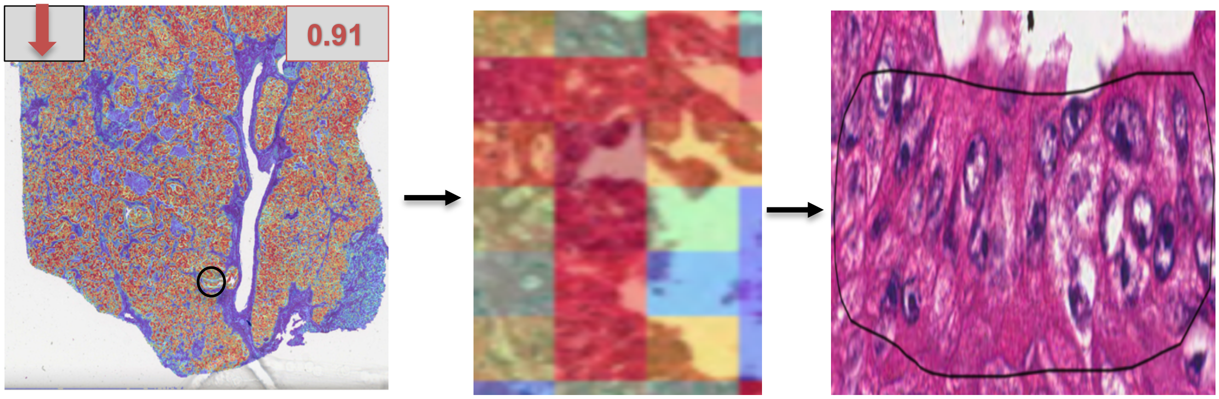


**Supplementary Figure S5**

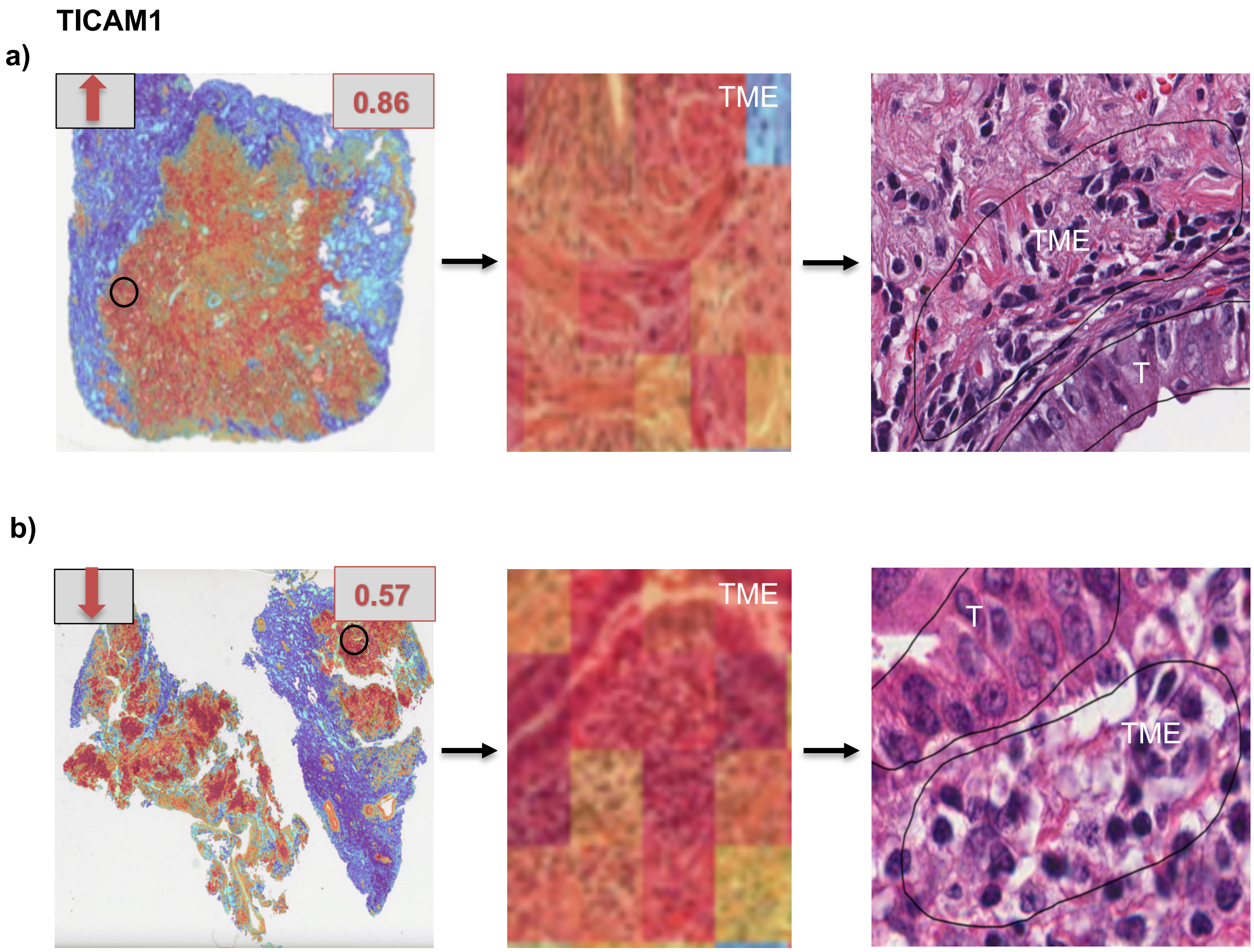


**Supplementary Figure S6**

**SLC47A1**

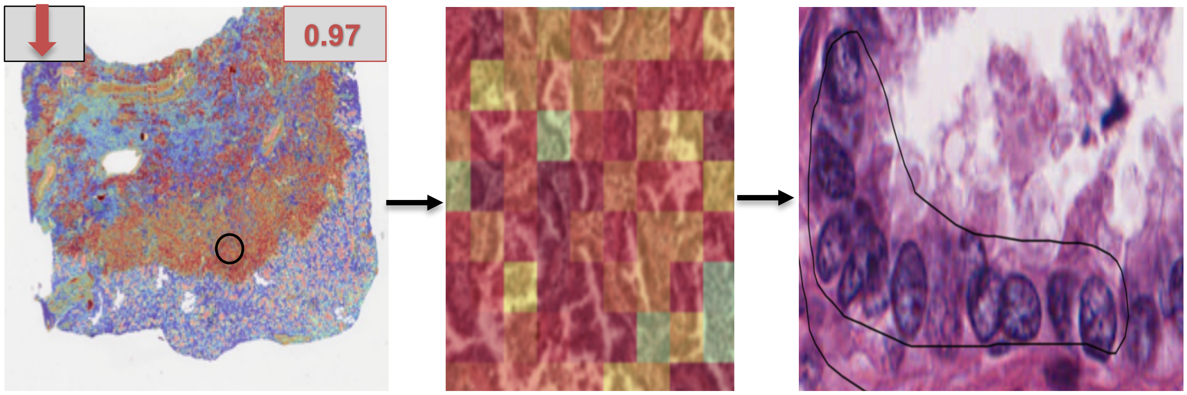


**Supplementary Figure S7**

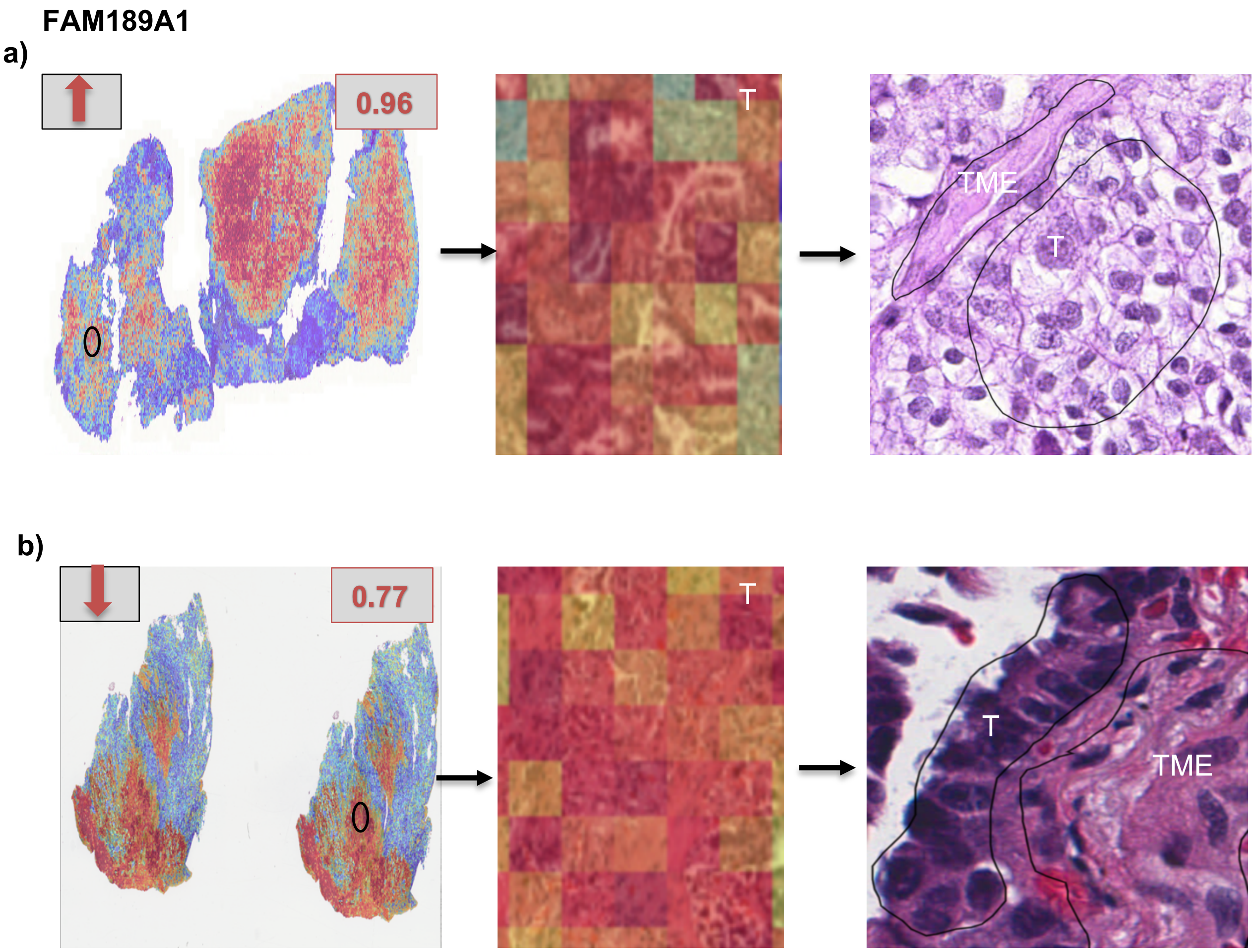


**Supplementary Figure S8**

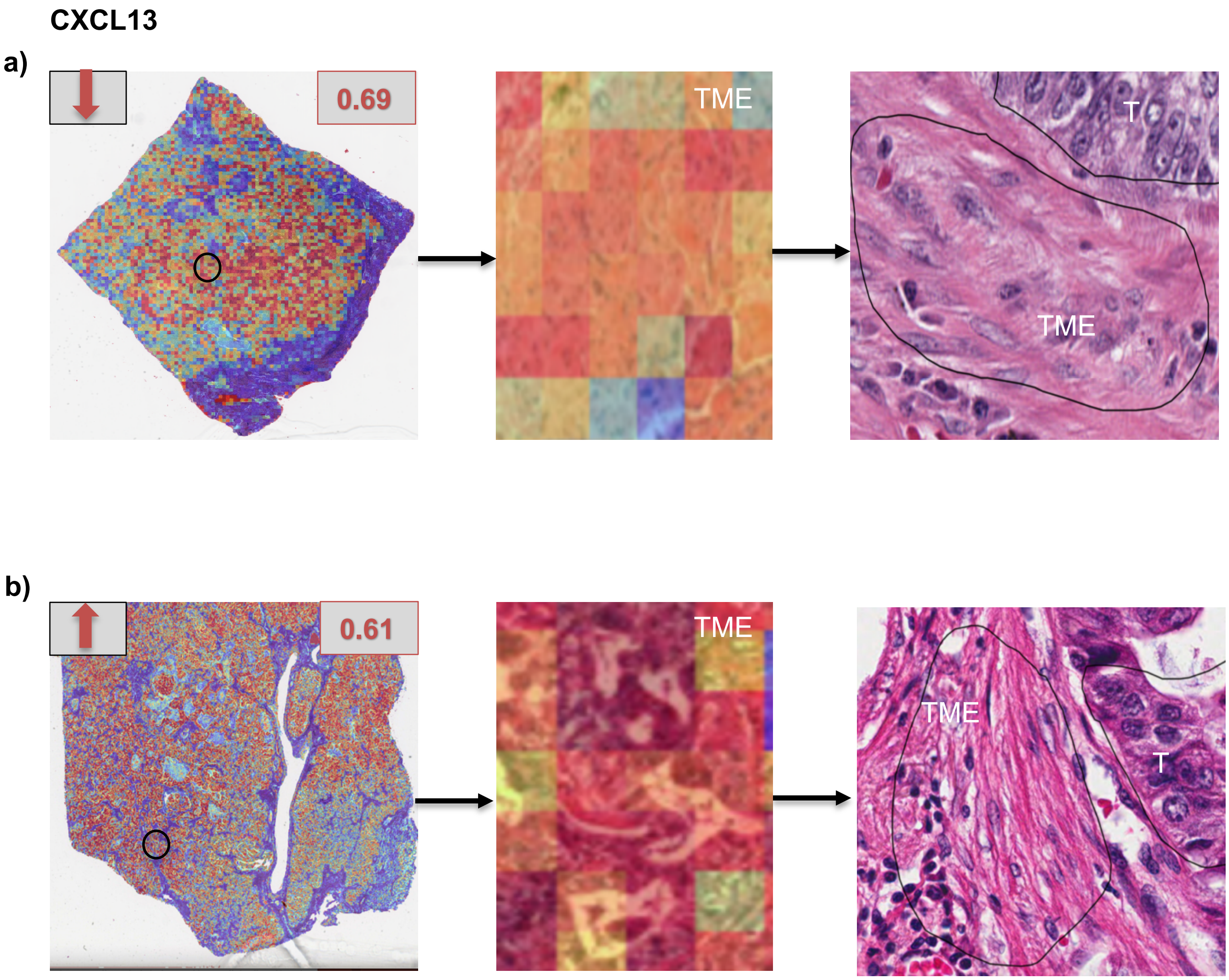


Supplementary Figure S9

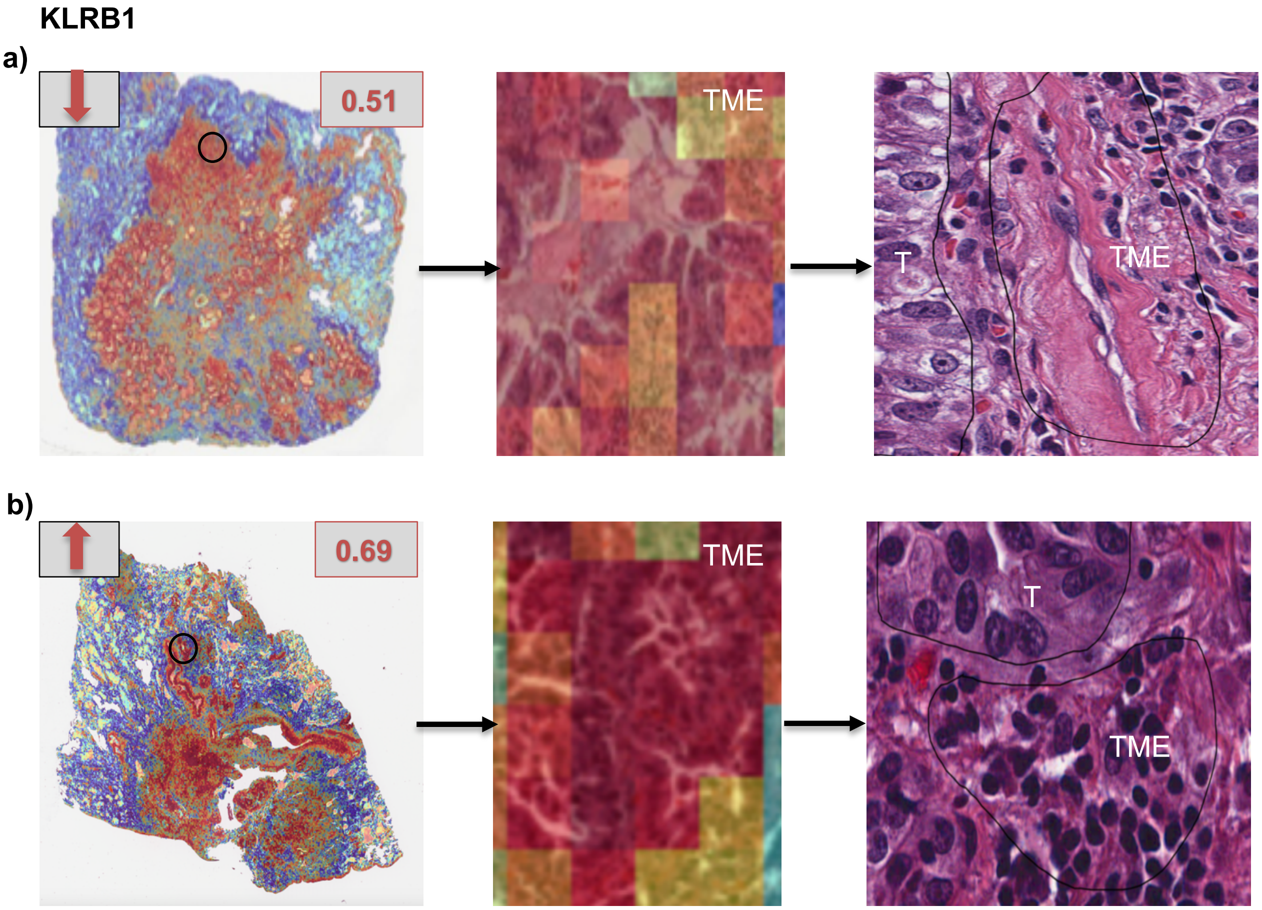


**Supplementary Figure S10**

**CDH3**

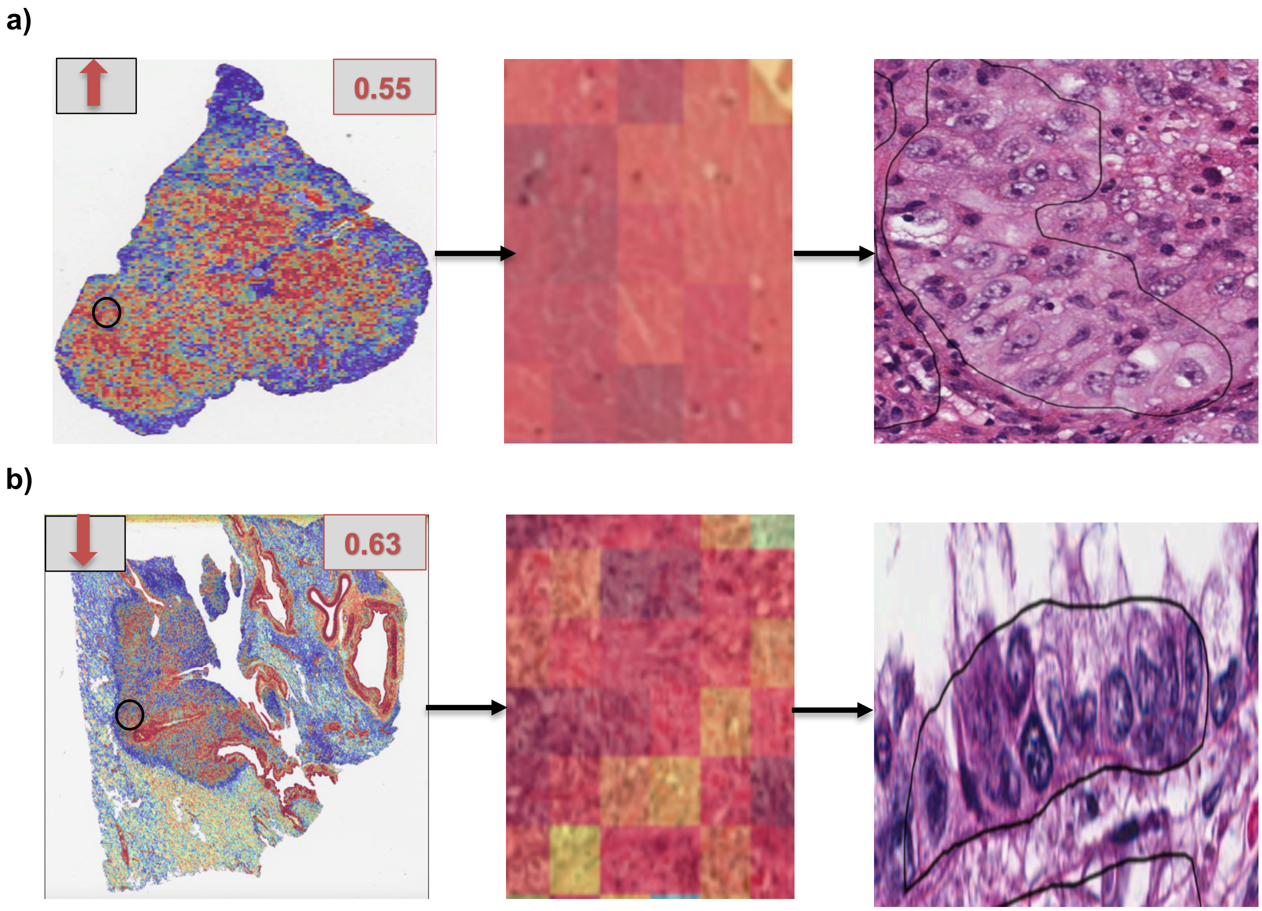


**Supplementary Figure S11**

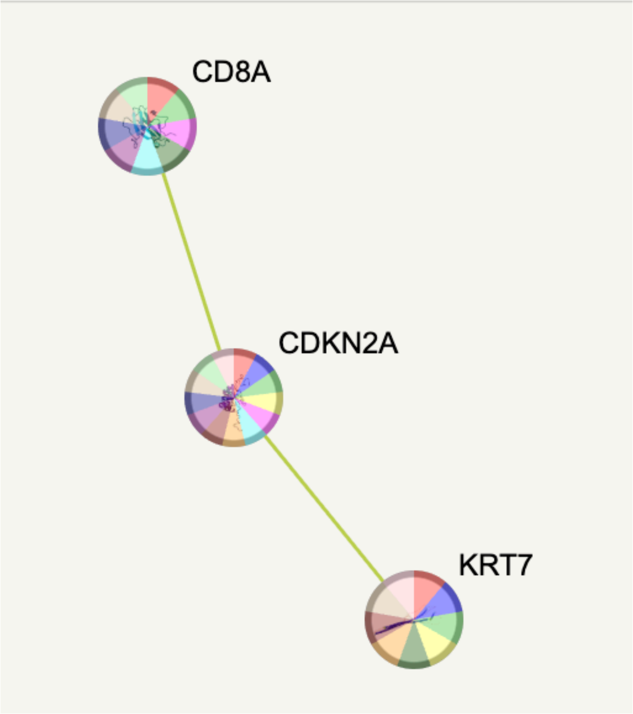


**Disease-gene Associations (DISEASES)**

| disease | description | count in network | strength | signal | false discovery rate |
|---|---|---|---|---|---|
| DOID:10155 | Intestinal cancer | 3 of 59 | 2.52 | 3.11 | 0.00014 |
| DOID:1294 | Vulva carcinoma | 2 of 5 | 3.42 | 2.87 | 0.00046 |
| DOID:3996 | Urinary system cancer | 3 of 89 | 2.35 | 2.85 | 0.00023 |
| DOID:403 | Mouth disease | 3 of 130 | 2.18 | 2.53 | 0.00046 |
| DOID:8618 | Oral cavity cancer | 2 of 18 | 2.86 | 2.39 | 0.0015 |
| DOID:10854 | Salivary gland disease | 2 of 18 | 2.86 | 2.39 | 0.0015 |
| DOID:9952 | Acute lymphoblastic leukemia | 2 of 26 | 2.7 | 2.23 | 0.0022 |
| DOID:4450 | Renal cell carcinoma | 2 of 26 | 2.7 | 2.23 | 0.0022 |
| DOID:3905 | Lung carcinoma | 2 of 36 | 2.56 | 2.08 | 0.0031 |
| DOID:3451 | Skin carcinoma | 2 of 39 | 2.53 | 2.05 | 0.0033 |
| DOID:11054 | Urinary bladder cancer | 2 of 38 | 2.54 | 2.05 | 0.0033 |
| DOID:5672 | Large intestine cancer | 2 of 44 | 2.47 | 2.02 | 0.0035 |
| DOID:1909 | Melanoma | 2 of 46 | 2.46 | 2.01 | 0.0036 |
| DOID:305 | Carcinoma | 3 of 307 | 1.81 | 1.9 | 0.0016 |
| DOID:9500 | Leukocyte disease | 2 of 77 | 2.23 | 1.74 | 0.0068 |
| DOID:170 | Endocrine gland cancer | 2 of 93 | 2.15 | 1.62 | 0.0090 |
| DOID:37 | Skin disease | 3 of 518 | 1.58 | 1.5 | 0.0037 |
| DOID:299 | Adenocarcinoma | 2 of 119 | 2.04 | 1.46 | 0.0135 |
| DOID:934 | Viral infectious disease | 2 of 133 | 1.99 | 1.39 | 0.0160 |
| DOID:3118 | Hepatobiliary disease | 2 of 168 | 1.89 | 1.24 | 0.0239 |
| DOID:0060085 | Organ system benign neoplasm | 2 of 237 | 1.74 | 1.03 | 0.0417 |

*(less ...)*

**Supplementary Figure S12-a**

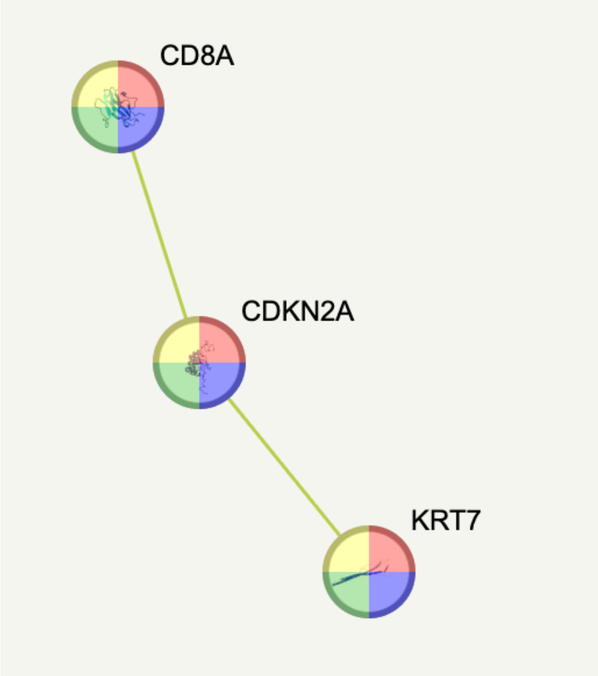


**Functional enrichments in your network**

explain columns

Reference Publications (PubMed)

| publication | (year) title | count in network | strength | signal | false discovery rate |
|---|---|---|---|---|---|
| PMID:39086316 | (2025) Tissue microarray validation in cervical carcinoma studies. ... | 3 of 5 | 3.6 | 3.13 | 0.00023 |
| PMID:36439155 | (2022) Case report: Local cryoablation combined with pembrolizu... | 3 of 11 | 3.25 | 2.77 | 0.00058 |
| PMID:35301248 | (2022) Clinical, radiological and pathological findings in patients wi... | 3 of 10 | 3.29 | 2.77 | 0.00058 |
| PMID:35024725 | (2021) Clinical and pathological aspects of condyloma acuminatu... | 3 of 13 | 3.18 | 2.77 | 0.00058 |
| PMID:34123576 | (2021) Prognostic image-based quantification of CD8CD103 T cell ... | 3 of 13 | 3.18 | 2.77 | 0.00058 |
| PMID:32300556 | (2020) Lack of Conventional Acinar Cells in Parotid Salivary Gland ... | 3 of 11 | 3.25 | 2.77 | 0.00058 |
| PMID:32095151 | (2020) Concurrence of Primary Cutaneous Extra Mammary Pagets ... | 3 of 11 | 3.25 | 2.77 | 0.00058 |
| PMID:27942391 | (2016) Metastatic basal cell carcinoma with amplification of PD-L1... | 3 of 13 | 3.18 | 2.77 | 0.00058 |
| PMID:26044703 | (2016) Absence of the intestinal microbiota exacerbates hepatobili... | 3 of 12 | 3.22 | 2.77 | 0.00058 |
| PMID:38611634 | (2024) Clinical, Epidemiological, Morphological, and Immunohistoc... | 3 of 16 | 3.09 | 2.76 | 0.00058 |
| PMID:34434351 | (2020) Local Lung Mass Masquerading a Very Aggressive Extraske... | 3 of 16 | 3.09 | 2.76 | 0.00058 |
| PMID:34368414 | (2021) Successful treatment of squamous cell carcinoma arising fr... | 3 of 15 | 3.12 | 2.76 | 0.00058 |
| PMID:30847303 | (2019) Putative Role of Circulating Human Papillomavirus DNA in t... | 3 of 14 | 3.15 | 2.76 | 0.00058 |
| PMID:37496666 | (2023) Case Report: Combined pembrolizumab, 5-fluorouracil, and ... | 3 of 17 | 3.06 | 2.75 | 0.00058 |
| PMID:35720298 | (2022) Upper Gastrointestinal Tract IrAEs: A Case Report About Sin... | 3 of 17 | 3.06 | 2.75 | 0.00058 |
| PMID:35763611 | (2022) The European Society of Gynaecological Oncology (ESGO), t... | 3 of 21 | 2.97 | 2.74 | 0.00058 |
| PMID:35728950 | (2022) The European Society of Gynaecological Oncology (ESGO), t... | 3 of 21 | 2.97 | 2.74 | 0.00058 |
| PMID:35004282 | (2021) Case Report: Tumor Microenvironment Characteristics in a ... | 3 of 23 | 2.93 | 2.74 | 0.00058 |
| PMID:33969717 | (2021) HTLV-1-Associated Lymphoma Presented as Massive Lymp... | 3 of 23 | 2.93 | 2.74 | 0.00058 |
| PMID:33866327 | (2021) Precision modeling of gall bladder cancer patients in mice b... | 3 of 22 | 2.95 | 2.74 | 0.00058 |
| PMID:29700411 | (2018) A dualistic model of primary anal canal adenocarcinoma wit... | 3 of 21 | 2.97 | 2.74 | 0.00058 |
| PMID:34832566 | (2021) Tissue Pathogens and Cancers: A Review of Commonly See... | 3 of 25 | 2.9 | 2.73 | 0.00058 |
| PMID:30718808 | (2019) Tumour-reactive T cell subsets in the microenvironment of ... | 3 of 24 | 2.91 | 2.73 | 0.00058 |
| PMID:27900363 | (2016) Integration of genomics and histology revises diagnosis an... | 3 of 24 | 2.91 | 2.73 | 0.00058 |
| PMID:21994886 | (2012) Tutorial review for understanding of cholangiopathy. | 3 of 25 | 2.9 | 2.73 | 0.00058 |
| PMID:20028992 | (2010) Useful immunohistochemical markers of tumor differentiati... | 3 of 26 | 2.88 | 2.73 | 0.00058 |
| PMID:35887164 | (2022) Are We Ready to Implement Molecular Subtyping of Bladder... | 3 of 27 | 2.86 | 2.71 | 0.00061 |
| PMID:37790755 | (2023) Cell-in-cell structure in cancer: evading strategies from anti-... | 3 of 29 | 2.83 | 2.68 | 0.00065 |
| PMID:36267983 | (2022) Update in collecting duct carcinoma: Current aspects of the ... | 3 of 29 | 2.83 | 2.68 | 0.00065 |
| PMID:36035166 | (2022) The combined prognostic model of copper-dependent to pr... | 3 of 30 | 2.82 | 2.68 | 0.00065 |
| PMID:35053578 | (2022) The Evolution of Ovarian Carcinoma Subclassification. | 3 of 28 | 2.85 | 2.68 | 0.00065 |
| PMID:34885178 | (2021) Identification of Survival and Therapeutic Response-Related... | 3 of 29 | 2.83 | 2.68 | 0.00065 |
| PMID:34113123 | (2021) Targeted-Gene Sequencing and Bioinformatics Analysis of ... | 3 of 28 | 2.85 | 2.68 | 0.00065 |
| PMID:33608032 | (2021) ARID1A genomic alterations driving microsatellite instability... | 3 of 28 | 2.85 | 2.68 | 0.00065 |
| PMID:33318584 | (2021) Massively parallel sequencing analysis of 68 gastric-type ce... | 3 of 29 | 2.83 | 2.68 | 0.00065 |
| PMID:32127601 | (2020) CTLA-4 blockade boosts the expansion of tumor-reactive C... | 3 of 30 | 2.82 | 2.68 | 0.00065 |
| PMID:24188515 | (2013) Lymphoepithelioma-like carcinoma of the stomach: a case r... | 3 of 28 | 2.85 | 2.68 | 0.00065 |
| PMID:37124415 | (2023) Alterations in histology of the aging salivary gland and corre... | 3 of 31 | 2.8 | 2.67 | 0.00065 |
| PMID:38299030 | (2024) VentX promotes tumor specific immunity and efficacy of im... | 3 of 33 | 2.78 | 2.62 | 0.00074 |
| PMID:37345129 | (2023) A Paradigm Shift in Primary Liver Cancer Therapy Utilizing G... | 3 of 33 | 2.78 | 2.62 | 0.00074 |
| PMID:38111767 | (2023) Metachronous primary esophageal squamous cell carcino... | 3 of 34 | 2.76 | 2.61 | 0.00076 |
| PMID:35008799 | (2021) High-Grade Cervical Intraepithelial Neoplasia (CIN) Associat... | 3 of 34 | 2.76 | 2.61 | 0.00076 |
| PMID:30598662 | (2018) How the BRAF V600E Mutation Defines a Distinct Subgroup ... | 3 of 34 | 2.76 | 2.61 | 0.00076 |
| PMID:22264787 | (2012) Bile acid and inflammation activate gastric cardia stem cell... | 3 of 37 | 2.73 | 2.54 | 0.00090 |
| PMID:37776168 | (2023) Patient-derived organoid culture in epithelial ovarian cancer... | 3 of 39 | 2.7 | 2.5 | 0.0010 |
| PMID:37727377 | (2023) Prognostic utility of TME-associated genes in pancreatic ca... | 3 of 42 | 2.67 | 2.5 | 0.0010 |
| PMID:37556095 | (2023) SEOM SOGUG clinical guideline for treatment of kidney can... | 3 of 42 | 2.67 | 2.5 | 0.0010 |
| PMID:36720497 | (2023) Immune and genomic biomarkers of immunotherapy respo... | 3 of 40 | 2.69 | 2.5 | 0.0010 |
| PMID:36304941 | (2022) Pulmonary Large Cell Neuroendocrine Carcinoma. | 3 of 39 | 2.7 | 2.5 | 0.0010 |
| PMID:35974300 | (2022) Construction of a novel model based on cell-in-cell-related g... | 3 of 40 | 2.69 | 2.5 | 0.0010 |
| PMID:35911327 | (2022) Synchronous Jejunal Sarcomatoid Carcinoma and Incidenta... | 3 of 39 | 2.7 | 2.5 | 0.0010 |
| PMID:33922215 | (2021) Intratumoral Cellular Heterogeneity: Implications for Drug R... | 3 of 40 | 2.69 | 2.5 | 0.0010 |
| PMID:32929114 | (2020) Integrative genomic analysis of salivary duct carcinoma. | 3 of 41 | 2.68 | 2.5 | 0.0010 |
| PMID:32015690 | (2020) The heterogeneous clinical and pathological landscapes of ... | 3 of 39 | 2.7 | 2.5 | 0.0010 |
| PMID:31786484 | (2020) American Registry of Pathology Expert Opinions: Evaluation... | 3 of 40 | 2.69 | 2.5 | 0.0010 |
| PMID:31455041 | (2019) BRAF Mutation in Colorectal Rhabdoid and Poorly Differenti... | 3 of 40 | 2.69 | 2.5 | 0.0010 |
| PMID:25934522 | (2015) Molecular staging of gynecological cancer: What is the futu... | 3 of 42 | 2.67 | 2.5 | 0.0010 |
| PMID:25114585 | (2010) Challenges in the development of future treatments for brea... | 3 of 40 | 2.69 | 2.5 | 0.0010 |
| PMID:37875494 | (2023) Nivolumab and ipilimumab in recurrent or refractory cancer ... | 3 of 43 | 2.66 | 2.49 | 0.0010 |
| PMID:35890003 | (2022) The Influence of Oncogenic Viruses in Renal Carcinogenesi... | 3 of 44 | 2.65 | 2.46 | 0.0011 |
| PMID:35886994 | (2022) MITFTFE Translocation Renal Cell Carcinomas: From Clinic... | 3 of 45 | 2.64 | 2.46 | 0.0011 |
| PMID:34436026 | (2021) Extramammary Pagets Disease: Diagnosis, Pathogenesis, a... | 3 of 47 | 2.62 | 2.42 | 0.0012 |
| PMID:30744199 | (2019) Non-Smoking-Associated Lung Cancer: A distinct Entity in T... | 3 of 46 | 2.63 | 2.42 | 0.0012 |
| PMID:29301290 | (2018) Non Melanoma Skin Cancer Pathogenesis Overview. | 3 of 46 | 2.63 | 2.42 | 0.0012 |
| PMID:37190324 | (2023) Cross-Dataset Single-Cell Analysis Identifies Temporal Alter... | 3 of 48 | 2.61 | 2.39 | 0.0013 |
| PMID:34522690 | (2021) An Overview of the Genomic Characterization of Hepatocell... | 3 of 48 | 2.61 | 2.39 | 0.0013 |
| PMID:37815881 | (2023) Identifying endoplasmic reticulum stress-related molecular ... | 3 of 50 | 2.6 | 2.36 | 0.0014 |
| PMID:36311715 | (2022) Case Report: Anlotinib combined with PD-1 inhibitor and se... | 3 of 51 | 2.59 | 2.34 | 0.0015 |
| PMID:30572031 | (2019) Best Practices Recommendations for Diagnostic Immunohi... | 3 of 51 | 2.59 | 2.34 | 0.0015 |
| PMID:33572888 | (2021) Immunohistochemical Biomarkers as a Surrogate of Molec... | 3 of 52 | 2.58 | 2.33 | 0.0015 |
| PMID:34232958 | (2021) Single-cell RNAseq and longitudinal proteomic analysis of a... | 3 of 53 | 2.57 | 2.31 | 0.0016 |
| PMID:33443130 | (2021) Genomic Alterations during the In Situ to Invasive Ductal Br... | 3 of 57 | 2.54 | 2.24 | 0.0019 |
| PMID:30050812 | (2018) Immunohistochemistry in Dermatopathology and its Releva... | 3 of 58 | 2.53 | 2.22 | 0.0020 |
| PMID:36419822 | (2022) Spatial intra-tumor heterogeneity is associated with survival... | 3 of 59 | 2.52 | 2.21 | 0.0021 |
| PMID:33930309 | (2021) Promotion of cholangiocarcinoma growth by diverse cancer... | 3 of 59 | 2.52 | 2.21 | 0.0021 |
| PMID:32922522 | (2020) Pulmonary sarcomatoid carcinoma: progress, treatment an... | 3 of 59 | 2.52 | 2.21 | 0.0021 |
| PMID:35860559 | (2022) Glia Maturation Factor beta as a Novel Independent Progno... | 3 of 62 | 2.5 | 2.19 | 0.0022 |
| PMID:33838281 | (2021) Understanding the cellular origin and progression of esopha... | 3 of 61 | 2.51 | 2.19 | 0.0022 |
| PMID:33005848 | (2020) Investigation of colorectal cancer in accordance with conse... | 3 of 61 | 2.51 | 2.19 | 0.0022 |
| PMID:34722263 | (2021) Single-Cell RNA Sequencing in Multiple Pathologic Types of ... | 3 of 65 | 2.48 | 2.14 | 0.0025 |
| PMID:35321284 | (2022) Pathological, molecular, and clinical characteristics of chola... | 3 of 67 | 2.47 | 2.11 | 0.0027 |
| PMID:33581928 | (2021) Molecular and immunological developments in placentas. | 3 of 69 | 2.46 | 2.08 | 0.0029 |
| PMID:37744328 | (2023) Isolation, culture, and delivery considerations for the use of ... | 3 of 72 | 2.44 | 2.03 | 0.0033 |
| PMID:35433771 | (2022) Up-to-Date Pathologic Classification and Molecular Charact... | 3 of 75 | 2.42 | 2.01 | 0.0035 |
| PMID:35205133 | (2022) Clinical and Molecular Characteristics of Rare Malignant Tu... | 3 of 74 | 2.43 | 2.01 | 0.0035 |
| PMID:34745998 | (2021) The Single-Cell Sequencing: A Dazzling Light Shining on the ... | 3 of 74 | 2.43 | 2.01 | 0.0035 |
| PMID:32560537 | (2020) Linking Cancer Stem Cell Plasticity to Therapeutic Resistan... | 3 of 74 | 2.43 | 2.01 | 0.0035 |
| PMID:35243422 | (2022) An omic and multidimensional spatial atlas from serial biop... | 3 of 76 | 2.41 | 2.0 | 0.0036 |
| PMID:40246777 | (2025) Therapeutic options for human papillomavirus-positive tons... | 2 of 2 | 3.82 | 1.98 | 0.0051 |
| PMID:40048885 | (2025) Development of a digital algorithm for assessing tumor-stro... | 2 of 2 | 3.82 | 1.98 | 0.0051 |
| PMID:39844467 | (2025) Deciphering the Differences Between Epstein-Barr Virus-Ass... | 2 of 2 | 3.82 | 1.98 | 0.0051 |
| PMID:39727619 | (2024) Digital Papillary Adenocarcinoma Is HPV-42-Associated and... | 2 of 2 | 3.82 | 1.98 | 0.0051 |
| PMID:39681999 | (2025) Nodal Yield From Neck Dissection Predicts the Anti-Tumor I... | 2 of 2 | 3.82 | 1.98 | 0.0051 |
| PMID:39614403 | (2024) Syringocystadenocarcinoma of the perianal region: a case r... | 2 of 2 | 3.82 | 1.98 | 0.0051 |
| PMID:39502617 | (2024) Prediction of Treatment Response Based on Nutritional Stat... | 2 of 2 | 3.82 | 1.98 | 0.0051 |
| PMID:38728457 | (2024) Identification and validation of a prognostic anoikis-related ... | 2 of 2 | 3.82 | 1.98 | 0.0051 |
| PMID:38321777 | (2024) Clinical significance of KRT7 in bladder cancer prognosis. | 2 of 2 | 3.82 | 1.98 | 0.0051 |
| PMID:38175704 | (2024) Homozygous Loss of CDKN2 in Primary Cutaneous CD8(+) ... | 2 of 2 | 3.82 | 1.98 | 0.0051 |
| PMID:37924082 | (2023) Explainable convolutional neural networks for assessing he... | 2 of 2 | 3.82 | 1.98 | 0.0051 |
| PMID:37795402 | (2023) Significance of neutrophil to lymphocyte ratio as a predictor... | 2 of 2 | 3.82 | 1.98 | 0.0051 |

(less ...)

**Supplementary Figure S12-b**

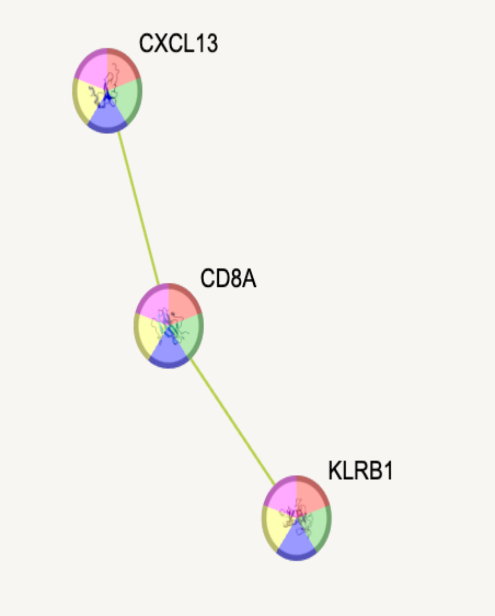


**Functional enrichments in your network**

*explain columns*

**Reference Publications (PubMed)**

| publication | (year) title | count in network | strength | signal | false discovery rate |
|---|---|---|---|---|---|
| PMID:37768011 | (2023) Single cell RNA sequencing sheds light on infiltrating T cells... | 3 of 7 | 3.45 | 2.85 | 0.00048 |
| PMID:40766325 | (2025) High-dimensional single-cell phenotyping unveils persistent ... | 3 of 14 | 3.15 | 2.44 | 0.0014 |
| PMID:34621785 | (2021) Fluorescent Multiplex Immunohistochemistry Coupled With ... | 3 of 17 | 3.06 | 2.41 | 0.0015 |
| PMID:28103284 | (2017) Gene Expression Profiling of Bronchoalveolar Lavage Cells ... | 3 of 17 | 3.06 | 2.41 | 0.0015 |
| PMID:36034981 | (2022) Single-cell immune ecosystem and metabolism reprogram... | 3 of 21 | 2.97 | 2.38 | 0.0016 |
| PMID:38200408 | (2024) Identification of CD8(+) T-cell exhaustion signatures for pro... | 3 of 22 | 2.95 | 2.37 | 0.0016 |
| PMID:37452311 | (2023) A necroptosis-related gene signature to predict prognosis a... | 3 of 23 | 2.93 | 2.37 | 0.0016 |
| PMID:38287014 | (2024) Semi-supervised integration of single-cell transcriptomics d... | 3 of 29 | 2.83 | 2.32 | 0.0018 |
| PMID:37294342 | (2023) Immunogenicity and efficacy of pembrolizumab and doxoru... | 3 of 28 | 2.85 | 2.32 | 0.0018 |
| PMID:36574662 | (2023) Identification of a unique subset of tissue-resident memory ... | 3 of 27 | 2.86 | 2.32 | 0.0018 |
| PMID:36505431 | (2022) In patients with primary Sjoegrens syndrome innate-like MA... | 3 of 30 | 2.82 | 2.32 | 0.0018 |
| PMID:34569985 | (2021) T cell dysfunction in glioblastoma: a barrier and an opportu... | 3 of 26 | 2.88 | 2.32 | 0.0018 |
| PMID:33717202 | (2021) Dysregulation of IL-17IL-22 Effector Functions in Blood and ... | 3 of 26 | 2.88 | 2.32 | 0.0018 |
| PMID:32973795 | (2020) Commentary: Group 3 innate lymphoid cells mediate early p... | 3 of 29 | 2.83 | 2.32 | 0.0018 |
| PMID:31683641 | (2019) Invariant NKT Cells and Rheumatic Disease: Focus on Prim... | 3 of 29 | 2.83 | 2.32 | 0.0018 |
| PMID:28480794 | (2018) Dimethyl fumarate induces changes in B- and T-lymphocyte ... | 3 of 29 | 2.83 | 2.32 | 0.0018 |
| PMID:22811678 | (2012) Relevance of regulatory T cell promotion of donor-specific t... | 3 of 28 | 2.85 | 2.32 | 0.0018 |
| PMID:38165493 | (2024) Exploring the relationship between immune heterogeneity c... | 3 of 32 | 2.79 | 2.31 | 0.0018 |
| PMID:36685593 | (2022) Disordered T cell-B cell interactions in autoantibody-positive... | 3 of 33 | 2.78 | 2.31 | 0.0018 |
| PMID:36526890 | (2022) The dynamics of gammadelta T cell responses in nonhuma... | 3 of 31 | 2.8 | 2.31 | 0.0018 |
| PMID:36186442 | (2022) miR-21-5pPRKCE axis implicated in immune infiltration and ... | 3 of 32 | 2.79 | 2.31 | 0.0018 |
| PMID:35508672 | (2022) Establishment of tissue-resident immune populations in the... | 3 of 34 | 2.76 | 2.31 | 0.0018 |
| PMID:35290245 | (2022) Single-cell transcriptomics reveals distinct effector profiles ... | 3 of 34 | 2.76 | 2.31 | 0.0018 |
| PMID:32580514 | (2020) Intratumoral Versus Circulating Lymphoid Cells as Predictiv... | 3 of 34 | 2.76 | 2.31 | 0.0018 |
| PMID:31602294 | (2019) Infection resisters: targets of new research for uncovering n... | 3 of 35 | 2.75 | 2.31 | 0.0018 |
| PMID:22277938 | (2012) A single intradermal injection of IFN-gamma induces an infl... | 3 of 32 | 2.79 | 2.31 | 0.0018 |
| PMID:37781383 | (2023) Multi-omics segregate different transcriptomic impacts of a... | 3 of 39 | 2.7 | 2.3 | 0.0018 |
| PMID:36901945 | (2023) Application of Machine Learning Models in Systemic Lupus ... | 3 of 39 | 2.7 | 2.3 | 0.0018 |
| PMID:36899020 | (2023) Immune subset-committed proliferating cells populate the h... | 3 of 38 | 2.71 | 2.3 | 0.0018 |
| PMID:33805904 | (2021) MAIT Cells: Partners or Enemies in Cancer Immunotherapy? | 3 of 39 | 2.7 | 2.3 | 0.0018 |
| PMID:33309739 | (2021) Single-cell RNA sequencing of psoriatic skin identifies path... | 3 of 40 | 2.69 | 2.3 | 0.0018 |
| PMID:33145436 | (2020) The identification and functional analysis of CD8+PD-1+CD... | 3 of 39 | 2.7 | 2.3 | 0.0018 |
| PMID:32582553 | (2020) The Cancer-Immune Set Point in Oesophageal Cancer. | 3 of 36 | 2.74 | 2.3 | 0.0018 |
| PMID:30770920 | (2021) T cells in primary Sjoegrens syndrome: targets for early inte... | 3 of 36 | 2.74 | 2.3 | 0.0018 |
| PMID:29247724 | (2018) Anti-HBV response to toll-like receptor 7 agonist GS-9620 is ... | 3 of 37 | 2.73 | 2.3 | 0.0018 |
| PMID:25595133 | (2015) Endothelial-binding, proinflammatory T cells identified by M... | 3 of 38 | 2.71 | 2.3 | 0.0018 |
| PMID:21959269 | (2011) MCAM-expressing CD4(+) T cells in peripheral blood secret... | 3 of 36 | 2.74 | 2.3 | 0.0018 |
| PMID:37544663 | (2023) Single-cell sequencing on CD8(+) TILs revealed the nature o... | 3 of 42 | 2.67 | 2.29 | 0.0018 |
| PMID:35508135 | (2022) Topologically associating domains are disrupted by evolutio... | 3 of 44 | 2.65 | 2.29 | 0.0018 |
| PMID:34939663 | (2022) The biological basis of disease recurrence in psoriasis: a hi... | 3 of 44 | 2.65 | 2.29 | 0.0018 |
| PMID:34745084 | (2021) The Tonsil Lymphocyte Landscape in Pediatric Tonsil Hyper... | 3 of 42 | 2.67 | 2.29 | 0.0018 |
| PMID:34495570 | (2021) Pathogenic T Cells in Celiac Disease Change Phenotype on ... | 3 of 43 | 2.66 | 2.29 | 0.0018 |

**Supplementary Figure S13**

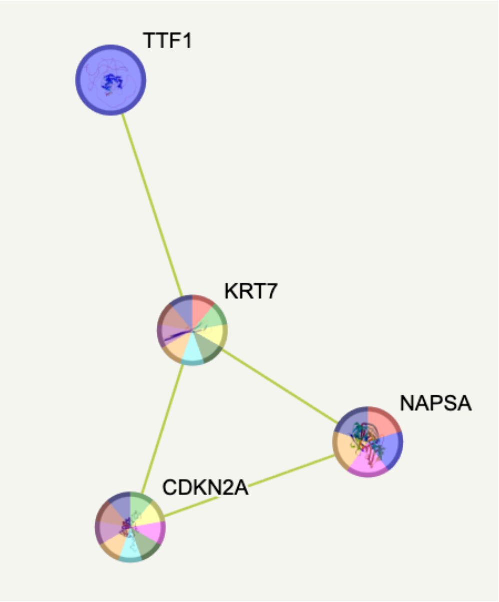


**Local Network Cluster (STRING)**

| cluster | description | count in network | strength | signal | false discovery rate |
|---|---|---|---|---|---|
| CL:20090 | Signet ring cell adenocarcinoma, and Hydrocele | 2 of 7 | 3.15 | 1.78 | 0.0051 |

**Reactome Pathways**

| pathway | description | count in network | strength | signal | false discovery rate |
|---|---|---|---|---|---|
| HSA-5683826 | Surfactant metabolism | 2 of 29 | 2.53 | 1.14 | 0.0328 |

**Disease-gene Associations (DISEASES)**

| disease | description | count in network | strength | signal | false discovery rate |
|---|---|---|---|---|---|
| DOID:1294 | Vulva carcinoma | 2 of 5 | 3.29 | 1.97 | 0.0030 |
| DOID:0050621 | Respiratory system benign neoplasm | 2 of 12 | 2.91 | 1.86 | 0.0039 |
| DOID:8618 | Oral cavity cancer | 2 of 18 | 2.74 | 1.8 | 0.0045 |
| DOID:3908 | Lung non-small cell carcinoma | 2 of 26 | 2.58 | 1.69 | 0.0060 |
| DOID:229 | Female reproductive system disease | 3 of 192 | 1.89 | 1.61 | 0.0039 |
| DOID:11054 | Urinary bladder cancer | 2 of 38 | 2.41 | 1.56 | 0.0085 |
| DOID:3451 | Skin carcinoma | 2 of 39 | 2.4 | 1.55 | 0.0085 |
| DOID:0060085 | Organ system benign neoplasm | 3 of 237 | 1.79 | 1.52 | 0.0046 |
| DOID:305 | Carcinoma | 3 of 307 | 1.68 | 1.35 | 0.0070 |
| DOID:10155 | Intestinal cancer | 2 of 59 | 2.22 | 1.35 | 0.0152 |
| DOID:28 | Endocrine system disease | 3 of 398 | 1.57 | 1.21 | 0.0101 |
| DOID:1100 | Ovarian disease | 2 of 109 | 1.96 | 1.05 | 0.0344 |
| DOID:299 | Adenocarcinoma | 2 of 119 | 1.92 | 1.01 | 0.0393 |
| DOID:0050686 | Organ system cancer | 3 of 757 | 1.29 | 0.79 | 0.0393 |

*(less ...)*

**Supplementary Figure S14-a**

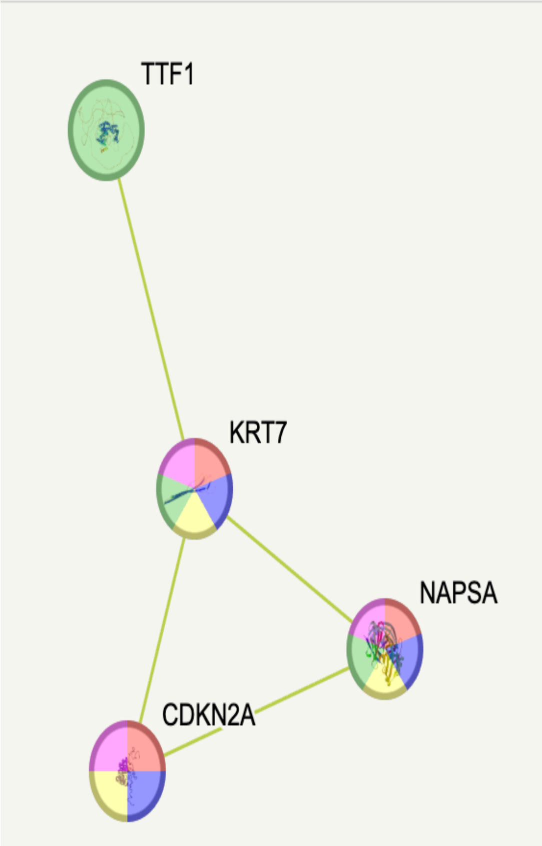


**Functional enrichments in your network**

explain columns

Reference Publications (PubMed)

| publication | (year) title | count in network | strength | signal | false discovery rate |
|---|---|---|---|---|---|
| PMID:37809183 | (2023) An Unusual Case of Nonbacterial Thrombotic Endocarditis ... | 3 of 3 | 3.69 | 2.73 | 0.00032 |
| PMID:38761046 | (2024) A Curious Case of Clear Cell Morphology in a Patient with L... | 3 of 7 | 3.32 | 2.47 | 0.00068 |
| PMID:32984377 | (2020) Expression of ALDH and SOX-2 in Pulmonary Sclerosing Pn... | 3 of 6 | 3.39 | 2.47 | 0.00068 |
| PMID:32303211 | (2020) Primary malignant melanoma of the lung: a case report and... | 3 of 6 | 3.39 | 2.47 | 0.00068 |
| PMID:29383242 | (2018) Metastatic squamous cell cancer of the lung presenting as ... | 3 of 7 | 3.32 | 2.47 | 0.00068 |
| PMID:23663245 | (2013) Lung cancer diagnosis on ovary mass: a case report. | 3 of 6 | 3.39 | 2.47 | 0.00068 |
| PMID:20823766 | (2011) Can we tell the site of origin of metastatic squamous cell ca... | 3 of 7 | 3.32 | 2.47 | 0.00068 |
| PMID:37675291 | (2023) Giant pulmonary sclerosing pneumocytoma with potentially... | 3 of 9 | 3.22 | 2.46 | 0.00068 |
| PMID:37194050 | (2023) Diffuse intrapulmonary mesothelioma mimicking pulmonar... | 3 of 10 | 3.17 | 2.46 | 0.00068 |
| PMID:35306333 | (2022) Clinicopathological feature of a resected large mixed squa... | 3 of 10 | 3.17 | 2.46 | 0.00068 |
| PMID:33134453 | (2020) Cutaneous metastasis from a vaginal squamous cell carcin... | 3 of 8 | 3.27 | 2.46 | 0.00068 |
| PMID:32215023 | (2020) Napsin-A Expression, a Reliable Immunohistochemical Mar... | 3 of 9 | 3.22 | 2.46 | 0.00068 |
| PMID:31567280 | (2020) Endometrial Gastric (Gastrointestinal)-type Mucinous Lesio... | 3 of 10 | 3.17 | 2.46 | 0.00068 |
| PMID:30845306 | (2018) Difficulties of clinical and histopathological diagnosis in adv... | 3 of 10 | 3.17 | 2.46 | 0.00068 |
| PMID:29266433 | (2018) Napsin A and WT 1 are useful immunohistochemical marke... | 3 of 9 | 3.22 | 2.46 | 0.00068 |
| PMID:28816950 | (2017) Pemetrexed-carboplatin with intercalated icotinib in the trea... | 3 of 8 | 3.27 | 2.46 | 0.00068 |
| PMID:28791269 | (2017) Primary mucinous adenocarcinoma of the vulva, intestinal t... | 3 of 9 | 3.22 | 2.46 | 0.00068 |
| PMID:27660815 | (2016) Vesical clear cell adenocarcinoma arising from endometrio... | 3 of 10 | 3.17 | 2.46 | 0.00068 |
| PMID:25408655 | (2014) A Poorly Differentiated Malignant Neoplasm Lacking Lung ... | 3 of 8 | 3.27 | 2.46 | 0.00068 |
| PMID:37592269 | (2023) Hereditary breast and ovarian cancer triggered by occult fall... | 3 of 11 | 3.13 | 2.45 | 0.00068 |
| PMID:37456237 | (2023) Case Report: Advanced pulmonary sarcomatoid carcinoma ... | 3 of 13 | 3.06 | 2.45 | 0.00068 |
| PMID:37109667 | (2023) A Case of Pneumothorax Ex Vacuo Associated with COVID-... | 3 of 11 | 3.13 | 2.45 | 0.00068 |
| PMID:36439155 | (2022) Case report: Local cryoablation combined with pembrolizu... | 3 of 11 | 3.13 | 2.45 | 0.00068 |
| PMID:36011067 | (2022) New Insights in the Diagnosis of Rare Adenocarcinoma Vari... | 3 of 11 | 3.13 | 2.45 | 0.00068 |
| PMID:35140975 | (2022) Giant basal cell carcinoma of anterior chest wall reveals me... | 3 of 12 | 3.09 | 2.45 | 0.00068 |
| PMID:34589958 | (2020) Novel MRPL13-ALK and PPP1CB-ALK Double Fusion As a P... | 3 of 13 | 3.06 | 2.45 | 0.00068 |
| PMID:33717766 | (2021) Cerebellopontine Angle Primary Choroid Plexus Carcinoma ... | 3 of 12 | 3.09 | 2.45 | 0.00068 |
| PMID:33392320 | (2020) Primary pulmonary malignant melanoma diagnosed with pe... | 3 of 12 | 3.09 | 2.45 | 0.00068 |
| PMID:33317482 | (2020) Adenocarcinoma arising from a foregut cyst of the diaphrag... | 3 of 11 | 3.13 | 2.45 | 0.00068 |
| PMID:33100952 | (2020) A rare case of gastric-type mucinous endocervical adenocar... | 3 of 13 | 3.06 | 2.45 | 0.00068 |
| PMID:31498173 | (2020) The Frequency and Prognostic Significance of the Histologi... | 3 of 11 | 3.13 | 2.45 | 0.00068 |
| PMID:31279344 | (2019) Primary endocervical gastric-type adenocarcinoma: a clinic... | 3 of 12 | 3.09 | 2.45 | 0.00068 |
| PMID:31049066 | (2019) Primary Clear Cell Adenocarcinoma of the Cervix: A Clinical ... | 3 of 13 | 3.06 | 2.45 | 0.00068 |
| PMID:29664741 | (2018) Primary Vaginal Gastric-type Adenocarcinoma and Vaginal ... | 3 of 11 | 3.13 | 2.45 | 0.00068 |
| PMID:38215117 | (2024) Whether specific genetic feature predicted immunotherapy ... | 3 of 17 | 2.94 | 2.44 | 0.00068 |
| PMID:38149229 | (2023) Metastatic Adenocarcinoma of Mandible with Unknown Pri... | 3 of 14 | 3.02 | 2.44 | 0.00068 |
| PMID:37946783 | (2023) Mucoepidermoid carcinoma of the lung with hemoptysis as ... | 3 of 15 | 2.99 | 2.44 | 0.00068 |
| PMID:37681079 | (2023) Rapid on-site evaluation of a solitary lung nodule in a patien... | 3 of 16 | 2.97 | 2.44 | 0.00068 |
| PMID:37663823 | (2023) Molecular pathology and clinical treatment of independent ... | 3 of 15 | 2.99 | 2.44 | 0.00068 |
| PMID:37533615 | (2023) p16 Immunohistochemical Expression in Nephrogenic Aden... | 3 of 14 | 3.02 | 2.44 | 0.00068 |
| PMID:37527987 | (2023) [Clinicopathological features of thyroid-like low-grade nasop... | 3 of 15 | 2.99 | 2.44 | 0.00068 |
| PMID:37288348 | (2023) Cracking the case: The diagnosis and treatment of a compl... | 3 of 16 | 2.97 | 2.44 | 0.00068 |
| PMID:37095820 | (2023) Nephrogenic Adenoma Arising From a Female Urethral Dive... | 3 of 14 | 3.02 | 2.44 | 0.00068 |
| PMID:36860892 | (2023) Rare Combined Small Cell Lung Carcinoma and Lung Squa... | 3 of 16 | 2.97 | 2.44 | 0.00068 |
| PMID:36159414 | (2022) Postoperative radiotherapy for thymus salivary gland carcin... | 3 of 15 | 2.99 | 2.44 | 0.00068 |
| PMID:35860058 | (2022) Clear cell carcinoma of the abdominal wall: A case report wi... | 3 of 14 | 3.02 | 2.44 | 0.00068 |
| PMID:35720298 | (2022) Upper Gastrointestinal Tract IrAEs: A Case Report About Sin... | 3 of 17 | 2.94 | 2.44 | 0.00068 |
| PMID:35382809 | (2022) Synchronous bilateral primary ovarian cancer with right end... | 3 of 14 | 3.02 | 2.44 | 0.00068 |
| PMID:35083213 | (2021) Case Report: Partial Response Following Nivolumab Plus D... | 3 of 16 | 2.97 | 2.44 | 0.00068 |
| PMID:34442412 | (2021) Tumour Genome Characterization of a Rare Case of Pulmon... | 3 of 17 | 2.94 | 2.44 | 0.00068 |
| PMID:33889618 | (2021) Gastrointestinal-type chemotherapy prolongs survival in an ... | 3 of 16 | 2.97 | 2.44 | 0.00068 |
| PMID:33680116 | (2021) Cervical malignant mixed mesonephric tumour: A case repo... | 3 of 15 | 2.99 | 2.44 | 0.00068 |
| PMID:33629595 | (2021) An Unexpected Case of Malignant Mesothelioma in a Young... | 3 of 16 | 2.97 | 2.44 | 0.00068 |
| PMID:33285759 | (2020) Breast metastasis from EGFRALK negative lung adenocarci... | 3 of 14 | 3.02 | 2.44 | 0.00068 |
| PMID:32316633 | (2020) Krukenberg Tumor in Association with Ureteral Stenosis Du... | 3 of 16 | 2.97 | 2.44 | 0.00068 |
| PMID:31375771 | (2019) Endometrial tumors with yolk sac tumor-like morphologic p... | 3 of 14 | 3.02 | 2.44 | 0.00068 |
| PMID:30634950 | (2019) Identification of a metastatic lung adenocarcinoma of the p... | 3 of 15 | 2.99 | 2.44 | 0.00068 |
| PMID:29338553 | (2018) Comprehensive Clinicopathologic and Updated Immunohist... | 3 of 14 | 3.02 | 2.44 | 0.00068 |
| PMID:37920167 | (2023) Metastasis of ovarian cancer to nasal skin and skin on the t... | 3 of 18 | 2.91 | 2.43 | 0.00068 |
| PMID:37818133 | (2023) Lymph node and bone metastasis of pulmonary intestinal a... | 3 of 19 | 2.89 | 2.43 | 0.00068 |
| PMID:36224612 | (2022) Identification of adenoid subtype characterized with immun... | 3 of 21 | 2.85 | 2.43 | 0.00068 |
| PMID:36207130 | (2022) Lorlatinib and compound mutations in ALK+ large-cell neur... | 3 of 19 | 2.89 | 2.43 | 0.00068 |
| PMID:35836509 | (2022) Primary pulmonary choriocarcinoma in male: report a case ... | 3 of 21 | 2.85 | 2.43 | 0.00068 |
| PMID:35110965 | (2022) Treatment Response to Immunotherapy After Crizotinib Res... | 3 of 21 | 2.85 | 2.43 | 0.00068 |
| PMID:33912440 | (2021) Case Report: Next-Generation Sequencing Reveals Tumor O... | 3 of 19 | 2.89 | 2.43 | 0.00068 |
| PMID:32118595 | (2020) HER2-positive Metastatic Melanoma: A Cautionary Tale! | 3 of 21 | 2.85 | 2.43 | 0.00068 |
| PMID:32022435 | (2020) Pulmonary sclerosing pneumocytoma: Cytomorphology and... | 3 of 21 | 2.85 | 2.43 | 0.00068 |
| PMID:31579423 | (2019) Multiple primary malignant neoplasms: A case report and lit... | 3 of 18 | 2.91 | 2.43 | 0.00068 |
| PMID:31174566 | (2019) Coexistence of endometrial mesonephric-like adenocarcino... | 3 of 18 | 2.91 | 2.43 | 0.00068 |
| PMID:30305059 | (2018) Combining genomic analyses with tumour-derived slice cult... | 3 of 20 | 2.87 | 2.43 | 0.00068 |
| PMID:29851704 | (2018) Diagnostic Algorithmic Proposal Based on Comprehensive I... | 3 of 18 | 2.91 | 2.43 | 0.00068 |
| PMID:29248205 | (2018) A guided tour of selected issues pertaining to metastatic ca... | 3 of 18 | 2.91 | 2.43 | 0.00068 |
| PMID:28494807 | (2017) Concordant clear cell mesonephric carcinoma of the bladde... | 3 of 20 | 2.87 | 2.43 | 0.00068 |
| PMID:38468567 | (2024) Primary cutaneous apocrine carcinoma with RARA::NPEPP... | 3 of 24 | 2.79 | 2.42 | 0.00068 |
| PMID:38124509 | (2023) Clinicopathological characteristics and treatment outcomes... | 3 of 24 | 2.79 | 2.42 | 0.00068 |
| PMID:38023125 | (2023) Synchronous mucinous metaplasia and neoplasia of the fe... | 3 of 22 | 2.83 | 2.42 | 0.00068 |
| PMID:36439460 | (2022) Histomorphological transformation from non-small cell lung... | 3 of 23 | 2.81 | 2.42 | 0.00068 |
| PMID:36004014 | (2022) KRAS G12C-Mutant Non-Small-Cell Lung Adenocarcinoma: ... | 3 of 22 | 2.83 | 2.42 | 0.00068 |
| PMID:35484217 | (2022) Intrahepatic cholangiocarcinoma hidden within cancer of un... | 3 of 22 | 2.83 | 2.42 | 0.00068 |
| PMID:35280487 | (2022) Prognostic value of an immunohistochemical signature in p... | 3 of 23 | 2.81 | 2.42 | 0.00068 |
| PMID:35204416 | (2022) Mesonephric-like Adenocarcinoma of the Ovary: Clinicopath... | 3 of 22 | 2.83 | 2.42 | 0.00068 |
| PMID:35004282 | (2021) Case Report: Tumor Microenvironment Characteristics in a ... | 3 of 23 | 2.81 | 2.42 | 0.00068 |
| PMID:34147056 | (2021) SMARCA4BRG1 protein-deficient thoracic tumors dictate re-... | 3 of 23 | 2.81 | 2.42 | 0.00068 |
| PMID:33321728 | (2020) Unusual Faces of Bladder Cancer. | 3 of 24 | 2.79 | 2.42 | 0.00068 |
| PMID:33198246 | (2020) Brain Metastasis from Unknown Primary Tumour: Moving fr... | 3 of 24 | 2.79 | 2.42 | 0.00068 |
| PMID:28451462 | (2017) Pulmonary adenocarcinoma with mucin production modula... | 3 of 24 | 2.79 | 2.42 | 0.00068 |
| PMID:26862903 | (2016) Distinguishing Lung Adenocarcinoma from Lung Squamous... | 3 of 24 | 2.79 | 2.42 | 0.00068 |
| PMID:26685087 | (2016) A Detailed Immunohistochemical Analysis of a Large Series... | 3 of 22 | 2.83 | 2.42 | 0.00068 |
| PMID:36253824 | (2022) Literature review of imaging, pathological diagnosis, and ou... | 3 of 25 | 2.77 | 2.41 | 0.00068 |
| PMID:33670088 | (2021) Mesonephric-Like Adenocarcinoma of the Endometrium: Di... | 3 of 26 | 2.75 | 2.41 | 0.00068 |
| PMID:29721309 | (2018) Unraveling endometriosis-associated ovarian carcinomas u... | 3 of 26 | 2.75 | 2.41 | 0.00068 |
| PMID:29101056 | (2018) Combined Small Cell Carcinoma of the Lung: Is It a Single E... | 3 of 26 | 2.75 | 2.41 | 0.00068 |
| PMID:28828031 | (2017) A diagnostically difficult case of a cellular pleural fluid: Mor... | 3 of 26 | 2.75 | 2.41 | 0.00068 |
| PMID:37510152 | (2023) CDX2- and PAX8-Expressing Subtypes in Female Urethral A... | 3 of 27 | 2.74 | 2.4 | 0.00069 |
| PMID:36385961 | (2022) Pulmonary Salivary Gland Tumor, Mucoepidermoid Carcino... | 3 of 27 | 2.74 | 2.4 | 0.00069 |
| PMID:34866915 | (2021) Favorable Response to Olaparib in a Patient with Cancer of ... | 3 of 27 | 2.74 | 2.4 | 0.00069 |
| PMID:29690599 | (2018) Clinicopathological Characteristics and Mutations Driving D... | 3 of 27 | 2.74 | 2.4 | 0.00069 |
| PMID:37394540 | (2023) Carcinoma of unknown primary (CUP): an update for histop... | 3 of 28 | 2.72 | 2.38 | 0.00074 |
| PMID:36998440 | (2023) Response to selpercatinib in a patient with RET fusion-positi... | 3 of 28 | 2.72 | 2.38 | 0.00074 |
| PMID:35053578 | (2022) The Evolution of Ovarian Carcinoma Subclassification. | 3 of 28 | 2.72 | 2.38 | 0.00074 |

(less ...)

**Supplementary Figure S14-b**
